# Transocular detection of premotor Parkinson’s disease via retinal capillary neurovascular coupling through functional OCT angiography

**DOI:** 10.1101/2024.08.04.606502

**Authors:** Kaiyuan Liu, Ruixue Wang, Longqian Huang, Huiying Zhang, Mengqin Gao, Bin Sun, Yizhou Tan, Juan Ye, Zhihua Ding, Ying Gu, Shaomin Zhang, Peng Li

**Author notes:** These authors contributed equally to this work (Kaiyuan Liu, Ruixue Wang). Corresponding author: Shaomin Zhang, and Peng Li, **Email:**, and. **Author Contributions:** K.L., R.W., S.Z., and P.L. designed research; K.L., R.W., L.H., H.Z., M.G., B.S., S.Z., and P.L. performed research; K.L., R.W., L.H., S.Z., and P.L. analyzed data; Y.T., J.Y., Z.D., Y.G., contributed new reagents/analytic tools; K.L., R.W., L.H., Y.T., J.Y., Z.D., Y.G., S.Z., and P.L. wrote the paper.

## Abstract

Early detection of Parkinson’s disease (PD) is essential for timely initiating neuroprotective interventions before significant loss of dopaminergic (DAergic) neurons, but faces challenges in accurately and noninvasively detecting subtle neuronal changes in the midbrain. Here, we propose a transocular functional imaging method termed fOCTA-rNVC, which detects premotor PD by measuring alterations in retinal neurovascular coupling (rNVC) at the capillary level using specialized functional OCT angiography (fOCTA). Our findings demonstrate that capillary rNVC is significantly attenuated and delayed due to concurrent retinal dopaminergic degeneration in premotor PD mice. Notably, this PD-related rNVC attenuation can be temporarily reversed in acute levodopa challenge. Utilizing the functional characteristics of capillary rNVC in PD, we achieved an impressive accuracy of ∼100% in detecting premotor PD mice even with only ∼14.1% loss of midbrain DAergic neurons, at which stage prompt treatment offered superior outcomes. In contrast, no significant changes were observed in retinal thickness or vasculature in the premotor PD mice. These findings suggest that fOCTA-rNVC is a promising noninvasive solution for accurately detecting premotor PD and guiding early interventions.

## Introduction

Parkinson’s disease (PD) is the second most prevalent neurodegenerative disorder worldwide (1). The main pathological mechanism underlying this condition is the gradual loss of midbrain dopaminergic (DAergic) neurons in the substantia nigra pars compacta (SNpc), which leads to dopamine (DA) depletion in the striatum and subsequent motor symptoms (2, 3). Currently, clinical diagnoses of PD are performed mainly by evaluating motor symptoms (4). However, by the time motor symptoms appear, a significant portion (70–80%) of DAergic neurons in the SNpc have already been lost, limiting the potential benefits of earlier neuroprotective therapies, which may slow, halt, or even reverse the progression of PD (5-8). Therefore, to detect premotor PD, it remains a challenge of non-invasively and accurately identifying subtle DAergic neuron degeneration within the deep brain.

Among efforts to detect PD at an early stage, numerous neuroimaging techniques have been developed to monitor the degeneration of DAergic neurons (9-11). Positron emission tomography (PET) and single-photon emission computed tomography (SPECT) target the presynaptic terminals of DAergic neurons, demonstrating high accuracy in identifying patients at risk of developing PD (9, 10). However, these approaches rely on radioactive isotopes, limiting their applicability in long-term monitoring of premotor PD. In contrast, the magnetic resonance imaging (MRI) is a noninvasive approach and can be used to examine the loss of nigral hyperintensity (11). Nevertheless, the sensitivity of MRI for detecting subtle changes that occur in the early stages of PD is very limited (12, 13). Moreover, these neuroimaging methods generally have high costs and limited accessibility, hindering their application in large-scale screening of PD.

The retina, an extension of the brain, could be examined to elucidate midbrain pathology in patients with PD (12, 14). Retinal DAergic degeneration is associated with the loss of midbrain DAergic neurons in PD patients (15), which may lead to retinal structural changes, such as reduced retinal thickness (16, 17) and decreased vessel density (18, 19). These abnormalities can be effectively investigated by optical coherence tomography (OCT) and OCT angiography (OCTA) but are found mainly in PD patients at advanced stages. Additionally, the results of a previous large cohort study suggested that the retinas of PD patients are only ∼1–2 µm thinner than those of healthy controls (16). Detecting such minor changes in thickness is challenging owing to the axial resolution of OCT, which is typically ∼4–7 µm (20, 21). Moreover, the PD-induced alterations of retinal vessel density are subtle (∼1%) and other diseases (e.g. diabetes) can also cause morphological changes in the retinal vasculature (18). These limitations, combined with individual variation, hinder the utilization of retinal structural biomarkers for the early screening of PD. Consequently, there is a need to explore additional retinal biomarkers that manifest in the early stages of PD and can be detected with greater precision.

In addition to structural changes, the degeneration of retinal DAergic neurons and the reduction in retinal DA levels are associated with functional changes in neuronal activity and retinal neurovascular coupling (rNVC). Electroretinography (ERG) has been used to measure the electrical neuronal response to light stimulation, revealing significantly attenuated and delayed electrical potentials in PD patients (22). Nevertheless, ERG requires corneal contact and is exceedingly time-consuming, limiting its clinical use in large populations (22, 23). In contrast, rNVC, which ensures that neuronal activity is matched by an appropriate change in blood flow to meet neuronal metabolic demands (24, 25), can be accurately assessed via noninvasive optical techniques. In functional OCTA (fOCTA), dynamic OCTA is employed to visualize retinal functional hyperaemia at the capillary level in response to synchronized flicker light stimulation (FLS) (26, 27), and the peak amplitude and time of the dynamic response are used as indices of rNVC function. Using fOCTA to measure rNVC as a potential biomarker of PD is a promising approach to address the unmet need for a noninvasive and accurate method for detecting premotor PD.

In this study, we present a novel optical functional approach, fOCTA-rNVC, for detecting premotor PD from the retina by using fOCTA to measure PD-related changes in rNVC at the capillary level (Fig. 1). First, we established a progressive PD mouse model characterized by a premotor stage with a slight loss of DAergic neurons in SNpc and retina and no evident motor deficits. Likely due to retinal DAergic degeneration, we found, for the first time, that functional rNVC is attenuated and delayed in premotor PD mice, whereas no significant change was found in the retinal structure, indicating the high sensitivity of rNVC as a functional biomarker for PD. Furthermore, we found that acute levodopa administration reversed PD-related rNVC attenuation in premotor PD mice, whereas no recovery was observed in ageing-related attenuation in aged mice without significant DAergic deficits, suggesting the high specificity of levodopa-recoverable rNVC in premotor PD. Finally, based on the levodopa recoverability of attenuated capillary rNVC, we achieved ∼100% accuracy in detecting premotor PD mice with ∼14.1% loss of midbrain DAergic neurons, underscoring the necessity of utilizing fOCTA to measure rNVC at the capillary level. Notably, the accurate detection of premotor PD facilitated prompt treatment with superior recovery. Therefore, the proposed fOCTA-rNVC is expected to address the unmet need for a noninvasive and accurate screening method for premotor PD patients, offering a cost-effective solution with superior accessibility and convenience for large-scale applications.

**Fig. 1.**
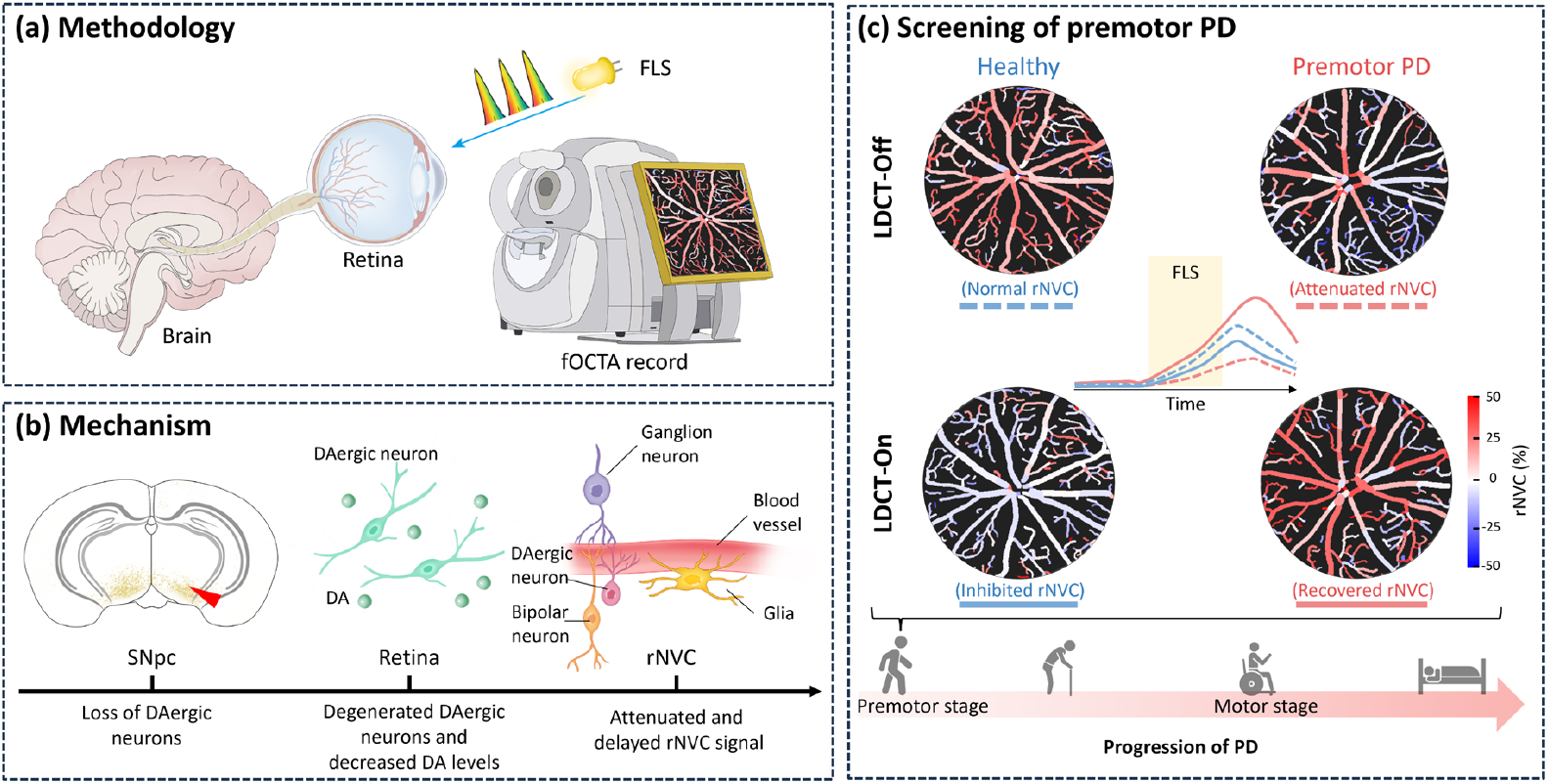
Schematic of the transocular detection of premotor PD by examining rNVC function with fOCTA. (a) In the proposed fOCTA-rNVC method, fOCTA is used to record PD-related changes in rNVC for the detection of premotor PD. In fOCTA, dynamic OCTA is employed to visualize retinal functional hyperaemia at the capillary level in response to synchronized FLS. (b) Association between PD and rNVC. Retinal DAergic degeneration is associated with the loss of SNpc DAergic neurons in PD, potentially leading to reduced DA levels in the retina and subsequently affecting neuronal activity and the associated rNVC. The red triangle indicates the obvious loss of DAergic neurons. (c) rNVC-based premotor PD detection. Premotor PD presents attenuated rNVC (red dashed curve, LDCT-Off) and levodopa-induced recovery (red solid curve, LDCT-On) which are different from healthy controls. FLS: flicker light stimulus; SNpc: substantia nigra pars compacta; DA: dopamine; rNVC: retinal neurovascular coupling; LDCT-Off: before levodopa challenge test; LDCT-On: during levodopa challenge test.

## Results

### SNpc and retinal DAergic degeneration in premotor PD mice

To determine whether changes in functional rNVC could be leveraged to identify PD before the onset of motor symptoms, we first sought to develop a premotor PD mouse model. To mimic the slight neurodegeneration associated with premotor PD, we unilaterally injected a low dose (0.35 μg) of 6-OHDA instead of the conventional dose (∼4.5 μg) reported in the literature (28), thereby inducing subtle but progressive damage to the DAergic neurons in the nigrostriatal system. We then sought to assess the suitability of our model by comparing physiological changes with those in sham mice, including assessments of neuronal degeneration and motor deficits.

One week after lesion induction, a minor decrease in the number of DAergic neurons in the ipsilateral SNpc (14.1%, *p* < 0.05, Figs. 2a-2b) and a slight decrease in the number of ipsilateral striatal fibres (26.3%, *p* < 0.01, Figs. 2c-2d) were observed in the PD group relative to those in the age-matched sham group. Such small-scale damage did not cause any significant behavioural deficits, as evidenced by the contralateral touches in the cylinder test (Fig. 2e) and the time spent on the pole test (Fig. 2f). As the disease progressed, the degree of degeneration in the nigrostriatal system increased, resulting in greater neuronal damage in the SNpc (58.2% neuronal loss, Fig. 2b) and striatum (60.5% fibre loss, Fig. 2d) at 3 weeks after lesion induction. This large-scale damage resulted in significant motor deficits in PD group with respect to the age-matched sham group (Figs. 2e-2f). Additionally, compared with the sham group, the number of turns in apomorphine-induced rotations by the PD mice substantially increased (*p* < 0.001, Fig. 2g), confirming extensive damage to the DAergic system in the PD model at 3 weeks after lesion induction. A detailed summary of the nigrostriatal pathology and behavioural impairments in the sham and PD groups is provided in Supplementary Table 1. Therefore, we successfully developed a premotor PD mouse model that showed mild nigrostriatal damage at 1 week after lesion induction, with significant motor symptoms appearing only in the third week.

**Fig. 2.**
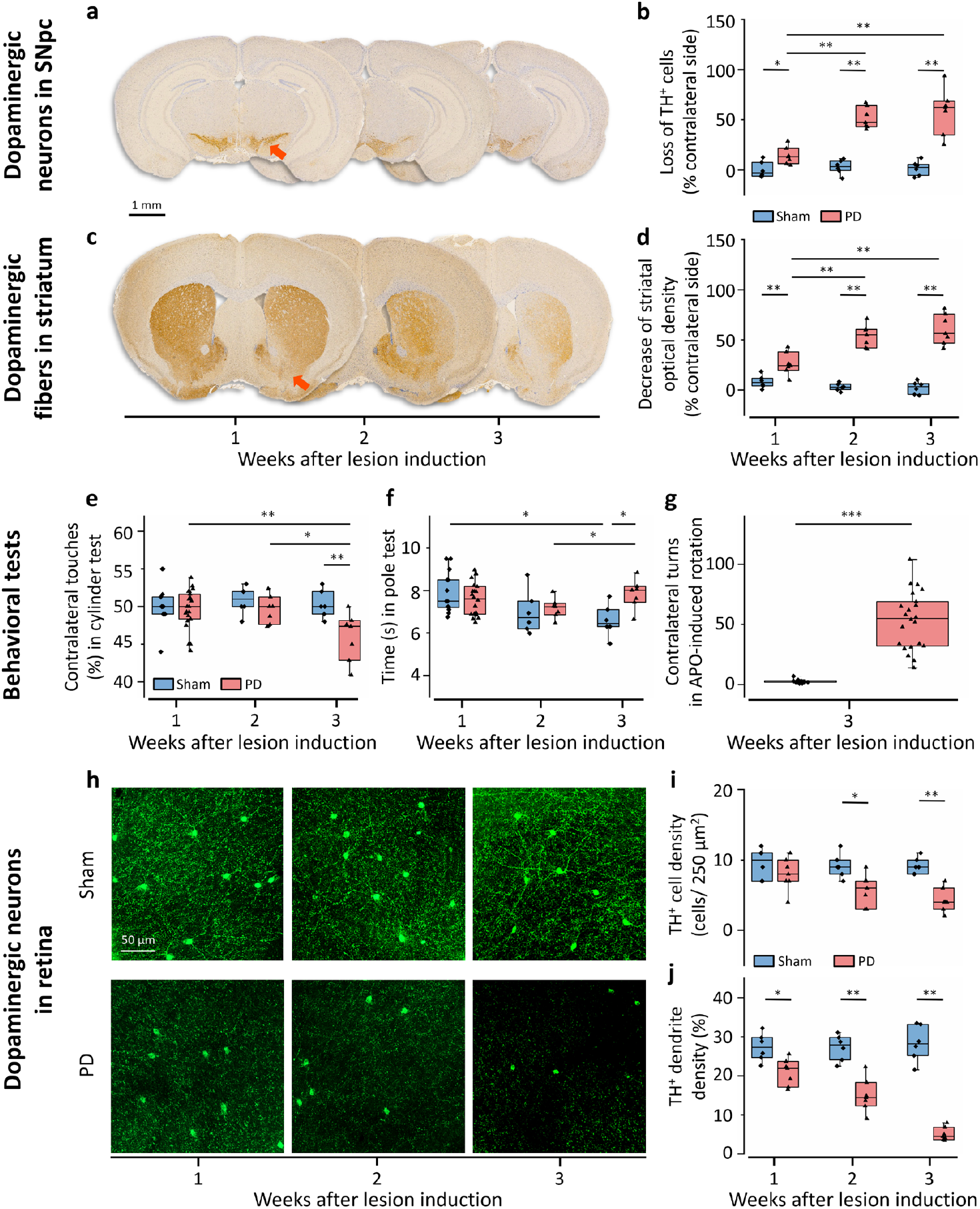
Retinal DAergic degeneration is accompanied by SNpc neuron loss in premotor PD mice. The PD mouse model was developed by unilaterally injecting low-dose 6-OHDA. Coronal brain sections stained for TH (yellow) showed the loss of DAergic neurons in the SNpc ipsilateral to the lesion side in PD mice (a), which was quantified by counting the number of SNpc TH^+^ cells (b) at 1 week (n = 6 Sham, 7 PD), 2 weeks (n = 6 Sham, 7 PD), and 3 weeks (n = 6 Sham, 7 PD) after lesion induction. The TH-stained (yellow) coronal brain sections also revealed DAergic degeneration in the striatum ipsilateral to the lesion side in PD mice (c), which was quantified by assessing the optical density in the striatum (d) at 1 week (n = 6 Sham, 7 PD), 2 weeks (n = 6 Sham, 7 PD), and 3 weeks (n = 6 Sham, 7 PD) after lesion induction. The red arrowheads denote areas in which there was an obvious reduction in the number of TH^+^ cells at 1 week after lesion induction. The cylinder test (e) and pole test (f) were used to evaluate behavioural deficits in sham and PD mice at 1 week (n = 13 sham, 22 PD), 2 weeks (n = 6 sham, 7 PD), and 3 weeks (n = 6 sham, 7 PD) after lesion induction. At 3 weeks after surgery, the mice in the sham group required less time than those at 1 week after surgery in the pole test (f), which may be attributed to the increased proficiency of the mice in the sham group following repeated testing. The APO-induced rotation test (g) was performed 3 weeks after surgery (n = 13 Sham, 22 PD) to verify the effectiveness of the 6-OHDA lesion induction protocol. (h) Whole-mount images of TH^+^ DAergic neuronal fibres and cell bodies (green) in the retinas of the sham and PD mice contralateral to the lesion side. Gradual degeneration of DAergic neurons was observed in the retinas of the mice in the PD group. The number of TH^+^ cell bodies per 250 µm2 (i) and dendrite density (j) at 1 week (n = 6 sham, 7 PD), 2 weeks (n = 6 sham, 7 PD), and 3 weeks (n = 6 sham, 7 PD) after surgery. TH: tyrosine hydroxylase, an enzyme involved in the synthesis of DA and used to identify DAergic cells; SNpc: substantia nigra pars compacta; APO: apomorphine; * *p* < 0.05, ** *p* < 0.01, and *** *p* < 0.001 for comparisons shown; one-tailed Mann–Whitney *U* test.

Similar to those in the nigrostriatal system, the DAergic neurons and dendrites in the retinas of PD mice gradually degenerated (Fig. 2h). Retinas from both the ipsilateral and contralateral lesion sides in the sham and PD groups were collected for immunohistochemical analysis. As early as 1 week after lesion induction, the density of retinal DAergic cell bodies was not significantly different between the contralateral retinas of PD and sham groups (Fig. 2i). However, the density of retinal DAergic dendrites in the contralateral retina significantly decreased to 20.9% in the PD group than that in the sham group (27.3%, *p* < 0.05, Fig. 2j). As retinal degeneration progressed, at 3 weeks after lesion induction, the cell body density and dendrite density in the contralateral retina of PD group further decreased to 4.3 cells/250 μm^2^ (*p* < 0.01) and 5.1% (*p* < 0.01), respectively. Notably, the densities of the cell bodies and dendrites in the ipsilateral retinas in the sham and PD groups and contralateral retina in the PD group did not significantly differ (Supplementary Table 1). In addition, no significant differences were detected in the densities of retinal endothelial cells and pericytes, the number of TUNEL^+^ neurons, or inducible nitric oxide synthase expression levels in the retinas between the contralateral PD group and the other three groups (Supplementary Fig. 1). Thus, SNpc neuron loss may lead to retinal DAergic degeneration in premotor PD. As there were no significant pathological differences among the retinas of the contralateral sham, ipsilateral sham, and ipsilateral PD groups, only the results of the analyses between the retinas in the contralateral PD and contralateral sham groups are presented in the following sections.

Our results indicate that the administration of low-dose 6-OHDA can be used to establish a progressive PD mouse model with a premotor stage characterized by 1) a slight reduction (∼14.1%) in the number of DAergic neurons in the SNpc; 2) the absence of evident motor deficits; and 3) a small but significant reduction in the number of DAergic dendrites in the retina.

### Attenuation and delay of functional rNVC in premotor PD

Given the intricate interplay between DA and NVC (29), we next sought to explore whether the subtle degeneration of DAergic neurons found in the retinas of premotor PD mice could affect functional rNVC, despite the lack of significant structural alterations in retinal thickness (Supplementary Fig. 2c). Specifically, we used fOCTA to assess the functional rNVC of premotor PD mice at 1-week post-lesion by assessing retinal functional hyperaemia in response to a given intensity of FLS (white light, 10 Hz, 50% duty ratio, 1000 lux, 30 s duration). To avoid the influence of motion-induced degradation of image quality and spatial misalignment during dynamic imaging, we developed techniques including temporal synchronization to control stimulus dosage, real-time display for monitoring imaging quality, and vascular skeleton-based registration algorithm to achieve motion alignment without losing vasodynamic information. Before FLS (baseline), the retinal trilaminar vascular network (the superficial (SCP), intermediate (ICP), and deep capillary plexuses (DCP), as shown in Fig. 3a) did not visibly differ between the sham and PD groups. Furthermore, no significant differences in the quantitative indices, including the vessel calibre (VC), vessel density (VD), vessel skeleton density, vessel complexity, flow velocity (FV), and retinal blood flow (RBF), were found (Supplementary Fig. 2d). Thus, neither the structural indices of retinal thickness and vasculature nor the baseline blood flow are sufficiently sensitive to reflect the subtle impairment in premotor PD mice.

**Fig. 3.**
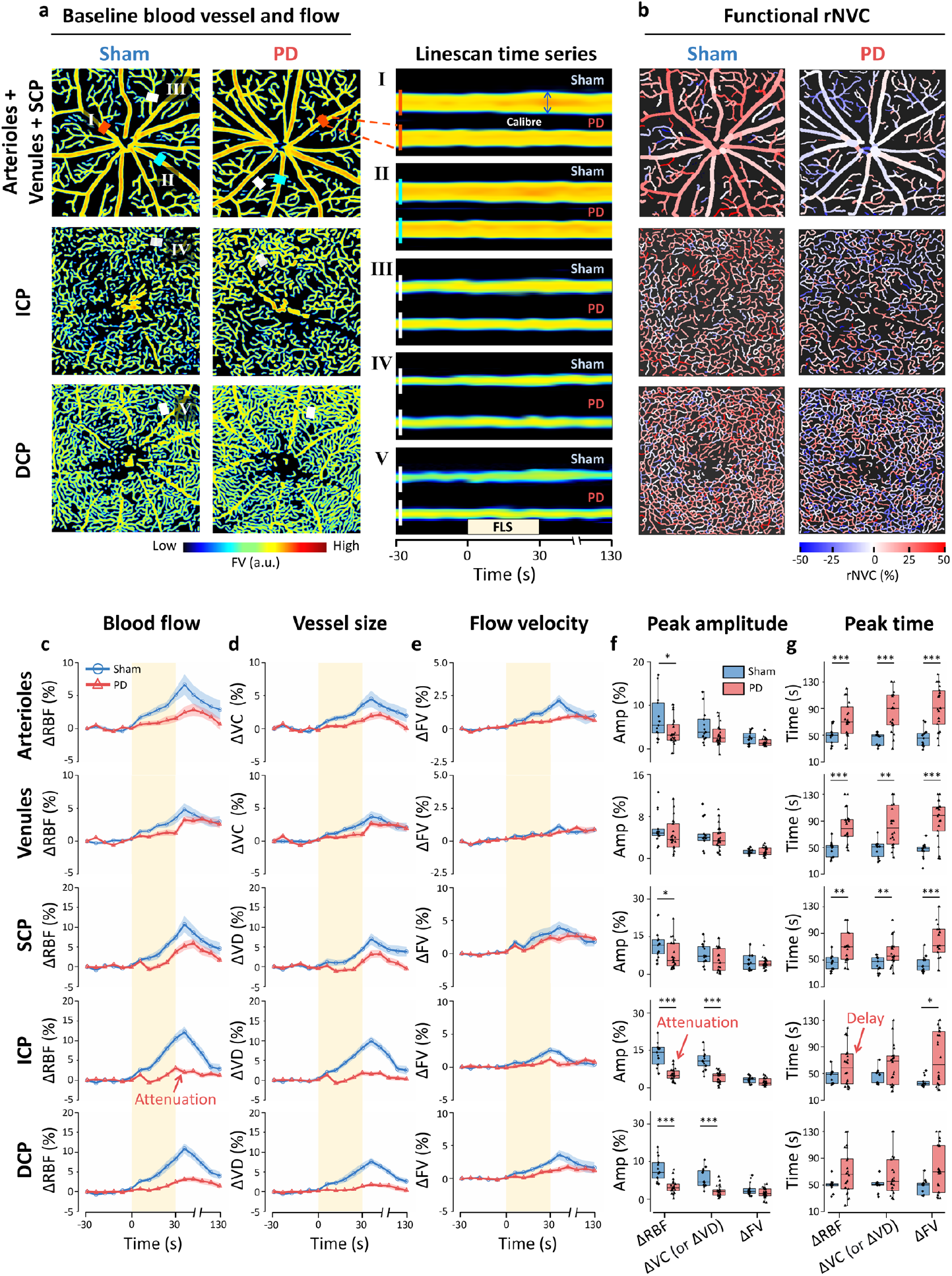
Functional rNVC signals are attenuated and delayed in premotor PD mice. (a) Baseline angiograms (encoded by the FV) of the trilaminar retinal vasculature (∼2 mm × 2 mm) in representative sham and PD retinas at 1 week after lesion induction. The insets (I-V) are linescan time series of functional hyperaemia before, during, and after FLS in a single arteriole, venule, and capillary (marked by the red, cyan, and white rectangles in the angiograms, respectively). Redder colours indicate higher FV values. (b) The corresponding functional rNVC map encoded by the peak amplitude of the ΔRBF time courses after FLS onset. The time courses of the blood flow (ΔRBF) (c), vessel size (ΔVC or ΔVD) (d), and blood flow velocity (ΔFV) (e) in the sham and PD mice. The yellow shaded regions indicate the period over which FLS was applied. Data are mean ± s.e.m.. The corresponding peak amplitude (f) and peak time (g) of functional rNVC signals in the arterioles, venules, SCP, ICP, and DCP are plotted. The retinas contralateral to the lesion side in the PD (n = 22) and age-matched sham (n = 13) groups at 1 week after lesion induction (no motor deficits present) were used for analysis. SCP: superficial capillary plexus; ICP: intermediate capillary plexus; DCP: deep capillary plexus; ΔRBF, ΔVC, ΔVD, and ΔFV: percentage changes in retinal blood flow, vessel calibre, vessel density, and flow velocity, respectively. FLS: flicker light stimulation. * *p* < 0.05, ** *p* < 0.01, and *** *p* < 0.001 for comparisons shown; one-tailed Mann–Whitney *U* test.

Unlike the retinal structural features, the functional indices of rNVC clearly differed between the PD and sham mice (Fig. 3b). rNVC function was assessed based on the FLS-evoked hyperaemic responses (en face angiographic video in Visualization 1 and Supplementary Fig. 3). Under FLS, remarkable hyperaemic responses, including vasodilation and increased blood flow, were observed in the linescan time series (the periods before, during and after FLS) of retinal vasculature in the sham mice but not in the PD mice (insets I-V in Fig. 3). To quantify the altered hyperaemic responses in the PD group, we calculated the percentage changes in the RBF (ΔRBF, Fig. 3c), vessel size (ΔVC or ΔVD, Fig. 3d) and FV (ΔFV, Fig. 3e) under FLS relative to the baseline values. Attenuated peak amplitudes and delayed peak times were observed in the time courses of the ΔRBF, ΔVC (or ΔVD), and ΔFV for the PD retinas. Compared with those in the sham retinas, significantly attenuated peak amplitudes were observed in the ΔRBF and ΔVD data obtained in the capillaries in the PD retinas (e.g., ICP in Fig. 3f: 5.2% vs. 13.4% of ΔRBF, *p* < 0.001). Furthermore, significantly longer delays (peak times) were observed for all three indices in the arterioles, venules, and SCP in the PD retinas than in the sham retinas (e.g., SCP in Fig. 3g: 69.3 s vs. 45.9 s for ΔRBF, *p* < 0.01). Since the ΔRBF reflects both the change in vessel size and flow velocity, we defined the peak amplitude of the ΔRBF time courses as the functional index of rNVC. Although obvious functional hyperaemia can be evoked with FLS, the responses presented a heterogeneity in both sham and PD mice, and vessels with positive (red) and negative (blue) rNVC index appeared simultaneously in the field of view (Fig. 3b). In particular, compared with those in the sham retinas, the incidence of arteriolar and venular dilation, which was defined as the percentage of dilated vessels (after FLS onset, the mean ΔVC > 0) among all vessels, significantly decreased in the PD retinas (Supplementary Figs. 4a-4b, arterioles: 68.6% vs. 89.9%, *p* < 0.01; venules: 70.5% vs. 90.9%, *p* < 0.01). Supplementary Table 2 presents a detailed summary of the functional hyperaemia observed in the sham and PD retinas with mean ± s.e.m. values.

Therefore, likely due to degeneration in the retinal DAergic system, functional rNVC signals are attenuated and delayed in premotor PD mice compared with those in healthy mice. However, no significant changes in retinal structure were observed, indicating the high sensitivity of functional rNVC as a biomarker for PD.

### Levodopa recoverability of PD-related rNVC attenuation

The above results indicate that functional rNVC is impaired in premotor PD mice. However, using only the observed attenuation and delay in the functional rNVC signal to identify PD has limitations, as rNVC could be affected by numerous neuronal and vascular conditions. For example, functional rNVC was also attenuated by ageing without DAergic deficits (Supplementary Fig. 5). Since PD is most common among elderly individuals, to improve the specificity of rNVC as a biomarker for premotor PD detection, we sought to explore whether levodopa, a precursor of DA used to treat PD in clinical settings (30), could reverse the attenuation in the functional rNVC signal in premotor PD and whether levodopa would have different effects on ageing-related attenuation.

To achieve this goal, we measured FLS-induced retinal functional hyperaemia before (LDCT-Off) and 1 h (LDCT-On) and 36 h (LDCT-Post) after the acute levodopa challenge test (LDCT) in PD, sham, and aged mice. The aged mice (44-week-old) were added to the experiment to demonstrate the specific rNVC response of PD mice via LDCT. The LDCT-Post measurements were obtained after the levodopa had fully metabolized (31). We found that the PD-related attenuation in the rNVC signals in the PD group was reversed at LDCT-On but returned at LDCT-Post (Fig. 4a). In contrast, the rNVC signals in the sham group at LDCT-On were even lower than those at LDCT-Off, and this attenuation was reversed at LDCT-Post (Fig. 4b). According to the quantitative plots of the functional rNVC signals, the attenuated hyperaemia was reversed in PD mice (ΔRBF in Fig. 4c; ΔVC, ΔVD, and ΔFV in Supplementary Figs. 6a and 6e), in contrast to the inhibited functional hyperaemia observed in response to FLS at LDCT-On in both the sham and aged retinas (ΔRBF in Figs. 4d-4e; ΔVC, ΔVD, and ΔFV in Supplementary Figs. 6b-6c and 6f-6g). Notably, an overshoot in the recovery of functional hyperaemia was observed in the PD group; that is, the ΔRBF signal not only recovered from its LDCT-Off value but also surpassed the LDCT-Off value in the sham mice (e.g., SCP in Fig. 3f: 21.3% vs. 12.4%, non-significance). However, we did not find any significant changes in peak time (Supplementary Fig. 7) or the incidence of arteriolar or venular dilation (Supplementary Figs. 4c-4d) among the PD, sham, and aged groups during LDCT. A detailed summary of the rNVC amplitudes (mean ± s.e.m.) measured during LDCT is provided in Supplementary Table 3.

**Fig. 4.**
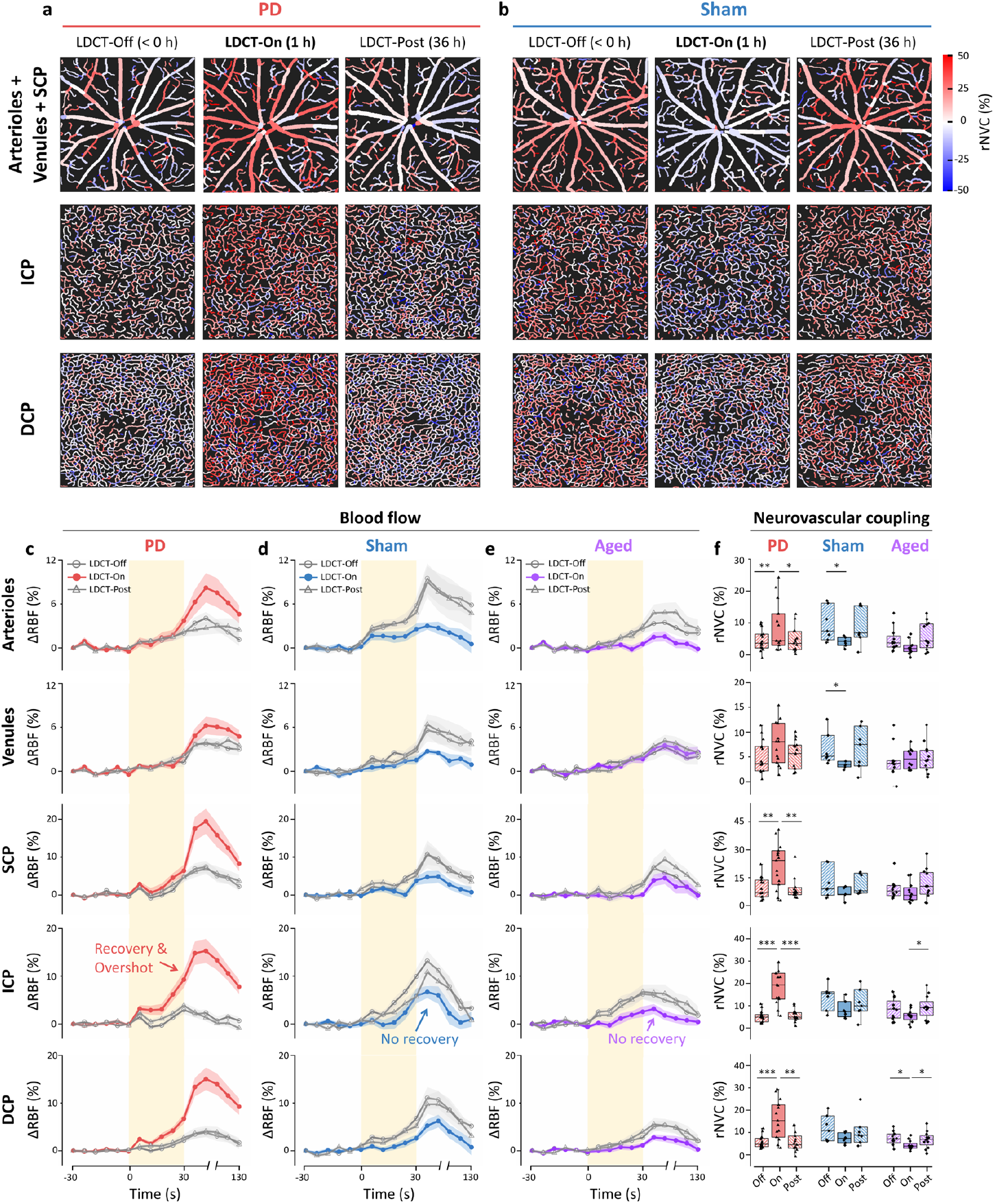
The PD-attenuated functional rNVC signal is recovered, whereas no recovery is observed in ageing-related attenuation during LDCT. Functional rNVC mapping (∼2 mm × 2 mm) of the PD (a) and sham (b) retinas before (LDCT-Off, < 0 h), during (LDCT-On, 1 h), and after LDCT (LDCT-Post, 36 h). The peak amplitude of the ΔRBF time courses after FLS onset was used as the rNVC index to encode the angiogram. The ΔRBF time courses in the PD (c), sham (d), and aged (e) retinas at LDCT-Off, LDCT-On, and LDCT-Post are shown. Data are mean ± s.e.m.. The yellow shaded regions indicate the period over which FLS was applied. (f) Box plots of the corresponding rNVC indices (peak amplitude of the ΔRBF time courses). The retinas contralateral to the lesion side in PD (n = 15, aged 12 weeks) and sham (n = 7, aged 12 weeks) mice at 1 week after lesion induction (no motor deficits present) and the retinas of aged mice (n = 12, aged 44 weeks) were used in the analysis. ΔRBF: percentage change in retinal blood flow; SCP: superficial capillary plexus; ICP: intermediate capillary plexus; DCP: deep capillary plexus; FLS: flicker light stimulation; LDCT: levodopa challenge test. * *p* < 0.05, ** *p* < 0.01, and *** *p* < 0.001 LDCT-On vs. LDCT-Off or LDCT-Post in the same group, Wilcoxon matched-pairs signed-rank test.

Therefore, the administration of levodopa has the potential to restore the attenuation in rNVC in premotor PD mice with deficits in the retinal DAergic system, whereas no recovery in the attenuation of rNVC is observed in mice without DAergic deficits. This finding indicates the high specificity of levodopa-recoverable rNVC as a functional biomarker for detecting premotor PD.

### Accurate detection of premotor PD facilitates prompt therapy

In the above sections, we demonstrated that the functional rNVC biomarker significantly differs between premotor PD and healthy mice, i.e. the PD shows attenuation and levodopa-induced recovery of rNVC. We then sought to explore whether premotor PD could be identified on the basis of 1) attenuation without levodopa administration or 2) both attenuation and levodopa-induced recovery and assessed the classification performance through receiver operating characteristic analysis.

Based solely on the attenuation of functional rNVC in PD, the functional index of ICP demonstrated a remarkable area under the receiver operating characteristic curve (AUROC) of up to 0.96 in effectively distinguishing premotor PD from Sham mice (Fig. 5a). In contrast, the AUROC values from the cylinder test (% contralateral touches), pole test (time taken to descend the pole), retinal thickness, and vascular morphology (baseline VC or VD values) did not exceed 0.62, highlighting the superior efficacy of functional rNVC as a biomarker for detecting premotor PD. However, the AUROC of the functional rNVC index in differentiating premotor PD mice and healthy aged mice was only 0.70 (Fig. 5b). This decrease in the AUROC is attributed to the significant attenuation observed in the rNVC signals in the aged group (Supplementary Fig. 5), which is similar to that observed in the PD group, potentially reducing the classification accuracy.

**Fig. 5.**
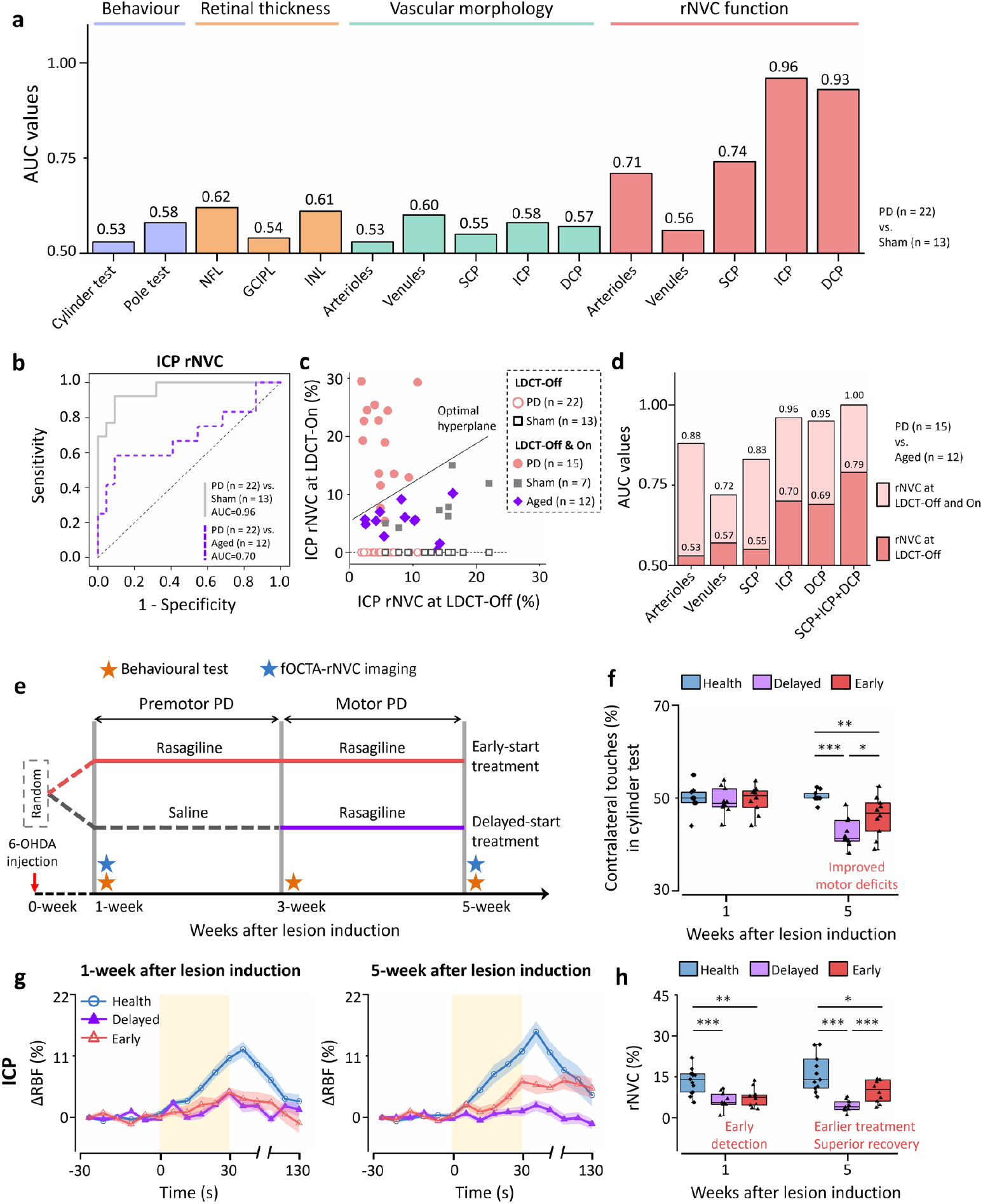
High accuracy of the rNVC test in detecting premotor PD facilitates prompt treatment with superior recovery. The rNVC indices are better in classifying premotor PD mice than behavioural test outcomes and retinal thickness and vascular morphology indices. (a) AUROC values for differentiating PD mice (n = 22) from age-matched sham (n = 13) mice on the basis of behavioural test outcomes and retinal thickness, vascular morphology (baseline VC or VD), and the rNVC index (LDCT-Off). NFL: nerve fibre layer; GCIPL: ganglion cell–inner plexiform layer; INL: inner nuclear layer. (b) Receiver operating characteristic curve analysis of the ability of the ICP rNVC index to distinguish PD (n = 22) from sham (n = 13) or aged (n = 12) mice at LDCT-Off. (c) Scatter plot of the ICP rNVC index at LDCT-Off (hollow symbols: PD, n = 22; sham, n = 13) and LDCT-Off & On (solid symbols: PD, n = 15; sham, n = 7; aged, n = 12). A support vector machine algorithm was applied to identify the optimal hyperplane for differentiating PD mice from both sham and aged mice based on the rNVC features at LDCT-Off and On (i.e. the solid symbols). (d) The AUROC values for discriminating PD mice (n = 15) from aged mice (n = 12) were calculated using one (LDCT-Off) or two (LDCT-Off and LDCT-On) rNVC features from the arterioles, venules, SCP, ICP, DCP, and the combination of SCP, ICP, and DCP. (e) Schematic illustration of the trial of treatment. First, the healthy mice aged 11 weeks were injected by 6-OHDA and randomly assigned to early-start (n = 10) and delayed-start (n = 10) treatment groups. The early-start group was treated with rasagiline since the observation of attenuated rNVC signals in premotor stage (from 1-week to 5-week after lesion induction), while the delayed-start group received saline for 2 weeks followed by rasagiline since the onset of motor symptoms (from 3-week to 5-week after lesion induction). Rasagiline is a potential neuroprotective drug in PD. The outcomes of the cylinder test (f) and the ΔRBF time courses of ICP (g) in the PD groups with early-start and delayed-start treatment and age-matched healthy sham mice at 1-week and 5-week after lesion induction. The yellow shaded regions in (g) indicate the period over which FLS was applied. Data are mean ± s.e.m.. (h) Box plots of the corresponding rNVC indices (peak amplitude of the ΔRBF time courses). AUROC: area under the receiver operating curve; SCP: superficial capillary plexus; ICP: intermediate capillary plexus; DCP: deep capillary plexus; ΔRBF: percentage change in retinal blood flow; LDCT: levodopa challenge test; FLS: flicker light stimulation. * *p* < 0.05, ** *p* < 0.01, and *** *p* < 0.001 for comparisons shown; one-tailed Mann–Whitney *U* test.

To further improve the accuracy, we used both attenuation and levodopa-induced recovery of rNVC for classification. The rNVC values at LDCT-Off (x-axis in Fig. 5c) and LDCT-On (y-axis in Fig. 5c) were both taken as features in a support vector machine model to generate an optimal hyperplane for differentiating PD mice from healthy control mice. Remarkably, we achieved even greater classification accuracy than using single rNVC feature at LDCT-Off (Fig. 5d, AUROC value increased from 0.70 to 0.96 in the ICP). Of note, by combining the rNVC features of SCP, ICP, and DCP at both LDCT-Off and LDCT-On, we further enhance the performance of PD detection to 1.00 AUROC (Fig. 5d, SCP+ICP+DCP). Therefore, compared with the behaviour test outcomes and retinal morphological features, the rNVC signals before and during LDCT can be used to effectively identify premotor PD. Notably, the highest accuracy was observed in the combination of trilaminar capillary plexus, demonstrating the importance of utilizing fOCTA to measure rNVC at the capillary level.

Enabled by the fOCTA-rNVC-based accurate detection of premotor PD, neuroprotective interventions could be initiated as early as possible. We conducted a trial of treatment (Fig. 5e) to demonstrate that prompt treatment leads to superior recovery compared to delayed treatment, and we aimed to explore whether the fOCTA-rNVC could be used to evaluate PD therapies. At 5-week after lesion induction, although the PD group treated before motor deficits still have motor deficits, it showed significant improvement than that treated after motor deficits (Fig. 5f and Supplementary Figs. 8a-8b). Notably, the fOCTA-measured rNVC function exhibited similar results in the behavioural tests, i.e. the rNVC indices of the PD treated before motor deficits showed superior recovery than that treated after motor deficits (Figs. 5g-5h and Supplementary Figs. 8c-8d).

Therefore, our results demonstrate that the fOCTA-rNVC can non-invasively and accurately detect premotor PD, which facilitates prompt treatment with superior recovery, and holds the potential to evaluate the efficacy of PD therapies.

## Discussion

Premotor PD detection allows for the implementation of neuroprotective therapies that could slow or stop disease progression (6, 8). However, accurately detecting the subtle degeneration of DAergic neurons at the early stage of PD remains a significant challenge. Here, we propose a novel proof of concept utilizing fOCTA-rNVC for premotor PD detection, which leverages fOCTA to accurately measure capillary rNVC as a robust biomarker. Through a customized fOCTA system, we demonstrated that functional rNVC is attenuated and delayed in premotor PD mice, and that the PD-attenuated functional rNVC is recoverable with levodopa. Additionally, on the basis of the levodopa recoverability of attenuated capillary rNVC, we achieved a remarkable accuracy of ∼100% in detecting premotor PD mice with ∼14.1% loss of midbrain DAergic neurons, at which stage prompt treatment offered superior prognosis. Overall, our findings support the potential of fOCTA-rNVC as a critical tool for the early detection and monitoring of PD.

The degeneration of retinal DAergic neurons in premotor PD mice is the pathological basis to establish our transocular detection approach. Our research indicates that the slight reduction in the number of SNpc DAergic neurons in premotor PD may be reflected in the retinal DAergic system, which may lead to a reduction in retinal DA levels, affecting the neuronal activity and the associated rNVC. Light-evoked neuronal activity can be recorded by ERG, and studies have shown attenuated and delayed electrical signals with levodopa-induced recovery in the retinas of PD patients (32, 33), which is similar to our rNVC signals observed at LDCT-Off and LDCT-On in the PD mice. These results may be attributed to that the DA levels are decreased in PD subjects (34), which markedly increased after the administration of levodopa (31). Although both neuronal activity and rNVC may be effective functional biomarkers for PD detection, to our knowledge, there have been no reports to date on the use of ERG in detecting premotor PD. In addition, conventional optical approaches have difficulty visualizing retinal neuronal activity in a label-free manner, and electrode-based methods have the potential to cause corneal or conjunctival abrasions. In contrast, the proposed rNVC biomarker can be extracted from retinal functional hyperaemia through fOCTA in a label-free, noncontact, and high-resolution approach. Therefore, rNVC dysregulation is a highly sensitive biomarker that is ideal for large-scale screening of premotor PD.

The levodopa-induced recoverability of rNVC demonstrates high specificity for detecting premotor PD. In clinical settings, LDCT is used to differentiate PD from other types of parkinsonism because PD patients typically show an improvement in motor symptoms when taking levodopa (30). Similarly, we combined rNVC examination with LDCT to show that attenuated rNVC could be improved after levodopa administration, suggesting that the observed attenuation in the rNVC signal may be attributed to DAergic degeneration. Furthermore, we confirmed that levodopa could not reverse the attenuated rNVC signal in aged mice, and no significant DAergic degeneration was observed in either the retina or the brain in these mice. Nevertheless, we will continue to evaluate the specificity of levodopa and other pharmacological drugs to rNVC in PD and other diseases. In future clinical practice, LDCT therapies could be applied to high-risk patients after initial fOCTA screening to mitigate potential side effects in low-risk populations.

In addition to the potential DA deficits in premotor PD, the observed rNVC overshoot under LDCT might be attributed to damage to DA receptors. DA receptors are crucial in DA-related signalling pathways, with D1-like receptors (D1/D5) primarily facilitating vasodilation and D2-like receptors (D2/D3/D4) promoting vasoconstriction (29). In healthy populations, the synergistic interplay between D1-like and D2-like receptors ensures adequate blood flow to meet neuronal demands (35); this balance is evidenced by the lack of significant differences in the total VC (or VD) and RBF values (baseline + FLS) between the LDCT-Off and LDCT-On conditions in both the sham and aged groups (Supplementary Figs. 9a-9c). Thus, the inhibited rNVC is most likely due to the increase in the baseline values due to levodopa-induced vasodilation, which subsequently limits the percentage changes in light-evoked hyperaemia in healthy mice. However, this balance might be disrupted in PD, as we observed a reversal from initial attenuation to a remarkable overshoot beyond normal levels after the administration of levodopa, which ultimately led to a significant increase in the total VC (or VD) at LDCT-On compared with that at LDCT-Off (Supplementary Figs. 9a, 9d and 9e). Our results indicate the presence of unregulated vasodilation in PD, likely due to the supersensitization of D1-like receptors and the impairment of D2-like receptors (36, 37).

We suggest the use of fOCTA rather than other retinal imaging techniques, such as dynamic vessel analysis (38) and ultrasound microscopy (39), to assess rNVC function because fOCTA can be used to measure capillary-level functional hyperaemia, which is essential for detecting premotor PD with high accuracy. Our results revealed that the functional imaging of the ICP resulted in better differentiation between the PD and sham groups compared to the major vessels, SCP, and DCP layers, with an AUROC value of 0.96. This remarkable performance may be attributed to the fact that the DAergic plexus is primarily distributed around the inner plexiform layer, resulting in the most severe functional damage to the ICP in PD retinas. In addition, we found that considering the ensemble of rNVC features in different vascular plexuses enhances the PD detection performance, indicating the potential of further improving the classification accuracy by leveraging rNVC features in multiple layers.

The fOCTA-rNVC-enabled premotor PD detection allows for prompt therapy implementation. Likely due to the neuroprotective effects of rasagiline on DAergic neurons (Supplementary Figs. 8e-8j), we demonstrated that the treatment since premotor stage leads to superior recovery in behavioural symptoms and rNVC function than that since motor stage. Thus, our methodology shows promise for objectively evaluating the efficacy of PD therapies. Although rasagiline could slow the degeneration of DAergic neurons, the early-treated PD group showed higher rNVC index at 5 weeks after lesion induction than that at 1 week. It may be attributed that the rasagiline could also increase DA levels by inhibiting DA metabolism (40, 41), leading to higher retinal DA concentration at 5 weeks post-lesion. Additionally, the 6-OHDA mouse model used in this study is widely employed in developing novel neuroprotective drugs for PD. Our fOCTA-rNVC facilitates quantitative monitoring of drug efficacy, thereby aiding drug discovery and development.

The main limitation of this study is the lack of clinical experiments involving premotor PD patients, primarily due to the requirement for long-term longitudinal monitoring to validate our method in clinical settings. Previous studies utilizing optical structural approaches have spanned over 3-7 years (16, 17). Unlike the structural method which can retrospectively analyze newly diagnosed PD patients using existing clinical datasets collected by conventional OCT setups, our functional method necessitates a new fOCTA system to collect rNVC data from scratch in PD patients since the premotor stage, resulting in prolonged clinical durations. Fortunately, our PD mouse model closely approximates the clinical setting for evaluating the accuracy of retina-based PD detection. In our study, the OCT retinal structural method demonstrated an AUROC accuracy of 62% in detecting premotor PD mice, which aligns well with the clinical result of 67% (17). Therefore, although clinical studies have not yet been conducted, we anticipate that our functional technique will also achieve significant improvements in detecting clinical PD patients similar to those observed in the PD mouse model. Currently, we are preparing clinical studies to evaluate our fOCTA-rNVC method in PD patients.

Overall, with a mouse model, we preliminarily demonstrated that the capillary rNVC signals measured by fOCTA are highly sensitive and specific for detecting premotor PD; moreover, the proposed approach is highly accessible and noninvasive. The noninvasive and accurate fOCTA-rNVC method has great potential for large-scale application in the screening of premotor PD patients, facilitating the initiation of neuroprotective interventions prior to the irreversible loss of DAergic neurons and potentially slowing PD progression. Furthermore, by assessing the recovery of impaired rNVC signals, fOCTA-rNVC can be utilized to evaluate the efficacy of PD treatments. Finally, the noninvasive, cost-effective, and portable attributes of our fOCTA system make it highly suitable for both community-based and at-home applications.

## Materials and Methods

### Animals

All surgical and experimental procedures conformed to the Guide for the Care and Use of Laboratory Animals (China Ministry of Health) and were approved by the Animal Care Committee of Zhejiang University (ZJU20220134). Male C57BL/6J mice (Zhejiang Medical Science Institute) were used in this study and divided into three groups: PD (n = 56, aged 11 weeks at lesion induction, body weight 23.7 ± 1.9 g), sham (n = 36, aged 11 weeks at saline injection, body weight 24.6 ± 2.2 g), and aged (n = 12, aged 44 weeks, body weight 33.5 ± 3.1 g). The animals were housed in an approved animal facility (ambient temperature of 22°C, relative humidity of 50%) under standard 12:12 h light:dark cycles, with food and water available ad libitum.

### PD model: 6-OHDA lesions

To mimic the premotor stage of PD, an animal model of PD was created in this study by intracerebrally injecting a low dose of 6-OHDA unilaterally into the medial forebrain bundle to create lesions in the mice. Briefly, 11-week-old mice were anaesthetized with isoflurane (4% for induction and 1.5–2% for maintenance). For the PD group, 6-OHDA solution (0.35 mg/ml) was then injected intracranially into the left medial forebrain bundle (centred 1.2 mm posterior and 1.2 mm lateral to the bregma and at a depth of 4.75 mm from the brain surface). A total of 1 µl of 6-OHDA was slowly infused at a rate of 0.5 µl/min using a glass capillary (42), which was left in the brain for an additional 3 min after the volume had been injected before being slowly withdrawn. The mice in the age-matched sham group were injected with 1 µl of saline solution at the same coordinates as the control mice. We verified the induction of the PD model using an apomorphine-induced rotational test at 3 or 5 weeks after lesion induction. Only mice in which hemiparkinsonian was successfully generated were included in the analysis of rNVC function.

### Behavioural tests

Here, we used drug-free evaluations, including the cylinder test and pole test, to assess motor symptoms in PD and age-matched sham mice after lesion induction. An apomorphine-induced rotational test was performed to verify the effectiveness of the 6-OHDA lesion induction method. All behavioural tests were performed by an observer blinded to experimental conditions.

The cylinder test is used to assess spontaneous forelimb asymmetry, reflecting the possibility that one of the forelimbs has poor function. Specifically, the mice were placed inside a transparent glass cylinder (inner diameter 10 cm, height 14 cm) and allowed to explore freely for 3 min while being recorded with a video camera. Two mirrors were placed accordingly so that all sides of the cylinder were visible to the camera. The film recording began once the mouse was placed inside with no habituation. In the 3-min recording period, each independent touch of the wall with the forelimbs ipsilateral and contralateral to the lesioned side was counted. Impairment of forelimb use was then calculated as the percentage of contralateral touches among all the touches with the following formula: (contralateral touches)/(ipsilateral touches + contralateral touches) ×100%.

The pole test is used to detect bradykinesia, one of the hallmark motor symptoms of PD, and evaluate motor coordination in PD mice. Each mouse was placed facing upwards atop a vertical wooden pole (diameter 1 cm, height 50 cm) within the home cage. The time the mouse took to descend the pole until it reached the floor of the home cage was recorded, with a maximum of 60 seconds. The mice were routinely pretrained for 3 or 4 days before the lesions were induced with the 6-OHDA protocol. During the pre– and postlesion tests, each mouse performed 5 successive trials with a 5 min intertrial interval. The average time across the 5 trials was recorded as the measured value.

In the apomorphine-induced rotational test, the mice were injected with 0.5 mg/kg apomorphine s.c. in the neck, placed in the recording chamber (a quiet room) and allowed to habituate to the environment for 10 min before the test began. The number of contralateral net rotations to the lesioned side (clockwise turns) within 60 min was recorded on video and then manually counted.

### Immunohistochemical staining

Brain staining was conducted as follows: first, the mice were anaesthetized with sodium pentobarbital and transcardially perfused with 0.9% saline, followed by paraformaldehyde (PFA) in phosphate-buffered saline (PBS). The brain was removed and postfixed for 12 h in 4% PFA at 4°C. Coronal slices of the striatum and SNpc were prepared using a cryostat (Leica, Germany). The free-floating slices were rinsed with PBS before quenching with 3% H_2_O_2_ to remove endogenous peroxidase for 10 min. After rinsing, the slices were permeabilized with 0.5% Triton X-100 for 20 min and incubated in 4% bovine serum albumin (Sigma Aldrich, USA) and 5% foetal calf serum for 1 h. The slices were then incubated overnight at 4°C with rabbit anti-mouse TH primary antibody (1:1000, Millipore, USA), followed by incubation for 20 min with a biotinylated secondary anti-rabbit antibody (1:200, LabVision, USA) and 10 min with a streptavidin– peroxidase complex at room temperature. Antigen visualization was performed using the chromogen 3,3’-diaminobenzidine. Finally, the slices were mounted on slides and dehydrated with an ethanol gradient and xylene. After TH staining, the stained slices were digitally scanned. The numbers of TH^+^ cells in the SNpc and the optical density of TH^+^ fibres in the striatum were quantified and are expressed as percentages relative to those in the intact hemisphere.

Retinal staining was conducted as follows: the eyes were enucleated and fixed in paraformaldehyde for 1 h at room temperature. The cornea and lens were removed, and the whole retina was dissected. Immunohistochemistry of retinal whole mounts and vertical sections was performed following previously described protocols (27). Briefly, retinal whole mounts were incubated with TH (1:200), IB4 (1:125), and NG2 (1:200, Abcam, UK) overnight at 4°C to immunolabel DAergic cells, endothelial cells, and pericytes, respectively. At the end of the incubation protocol, the sections were washed 3 times with PBS and then incubated with secondary antibodies (1:100, Thermo Fisher, USA) for 1 h. Retinal sections were deparaffinized and incubated with a primary antibody against inducible nitric oxide synthase (iNOS) (1:200, Proteintech, USA) overnight at 4°C to immunolabel iNOS. Then, secondary antibodies (1:100) were applied for 1 h. In addition, apoptotic neurons were detected via TUNEL, and the cell nuclei were stained with DAPI. Fluorescence images were obtained with a confocal microscope. The total number of DAergic and TUNEL^+^ cells was counted, and the densities of DAergic dendrites, endothelial cells, pericytes, and iNOS were calculated.

### Preparation for fOCTA imaging

The retinas contralateral to the lesioned side of the PD and sham mice and the bilateral retinas of the aged mice were imaged via fOCTA. All the experiments were conducted in a dark room, and all ambient light was blocked. Prior to the experiments, the mice were first dark-adapted for 1 h and then anaesthetized with 1% pentobarbital as previously described. After anaesthesia induction, the animal was immobilized in a laboratory-designed animal holder for fOCTA device alignment. The pupils were dilated with drops of phenylephrine hydrochloride and tropicamide, and then gel tears and contact lenses (Unicon, Japan) were applied to maintain a hydrated cornea and prevent cataract formation during the fOCTA imaging sessions. The animal’s body temperature was maintained at ∼36.5°C using a heating blanket (21). All fOCTA scans were conducted before noon to reduce the influence of diurnal variation.

### fOCTA system

The fOCTA system (Supplementary Fig. 10) used here was a custom-built prototype that is composed of two main modules: one module is used to provide the visual stimulus, and another module is used to monitor retinal hyperaemia (27). The visual stimulus module generates a diffuse flicker light with a white LED to increase the neuronal metabolic demand and evoke a haemodynamic response. The hyperaemia monitoring module is a time-lapse spectral-domain OCTA system (26) for rNVC signal collection. The system has an illumination spectrum covering the weak-absorption contrast region of water (λ_*central*_ = 840 nm; full width at half maximum of ∼100 nm). It operates at a 120 kHz axial sampling rate, with axial and lateral resolutions of ∼3 μm and ∼10 μm in the retina. The FLS was triggered during the imaging sequence by a circuit, ensuring synchronization between the visual stimulus and the OCTA recording. The total light power of the OCT and FLS on the pupil was ∼1 mW, which is within the safety level allowed by the American National Standards Institute (43).

### Stimulation protocol

The FLS pattern included a 30-s baseline period, a 30-s stimulation period, and a 100-s poststimulation period. During the stimulation period, the mean illuminance of the FLS on the cornea was 1000 lux (10 Hz, 50% duty ratio).

### Acute levodopa challenge test

One week after lesion induction, after the behavioural tests and fOCTA imaging at LDCT-Off, premotor PD (n = 15), sham (n = 7), and healthy aged (n = 12) mice were selected for LDCT. Levodopa was orally administered to the mice at a single dose of 20 mg/kg body weight. fOCTA imaging was conducted at 1 h (LDCT-On) and 36 h (LDCT-Post) after levodopa administration.

### Study design of treatment

After the injection of 6-OHDA, n = 20 PD mice (aged 11 weeks) were randomly assigned to early-start (n = 10) and delayed-start (n = 10) treatment groups. The early-start treatment group received a daily subcutaneous injection of rasagiline (2.5 mg/kg) as early as the observation of attenuated rNVC signals in premotor stage (from 1-week to 5-week after lesion induction). In contrast, the delayed-start treatment group received saline for 2 weeks followed by rasagiline since the onset of motor symptoms (from 3-week to 5-week after lesion induction). The fOCTA-rNVC imaging was conducted at 1-week and 5-week. The cylinder test and pole test were conducted at 1-week, 3-week, and 5-week after lesion induction. After the fOCTA imaging at 5-week post-lesion, the apomorphine-induced rotation test was performed.

### Data acquisition

During dynamic imaging, repeated volumetric raster scans (z-x-y) centred on the optic nerve head with a field of view of ∼2 mm × 2 mm (x-y) were collected at each timepoint. Each volumetric scan was acquired within 2 s and consisted of 256 axial profiles (x) to form a B-scan, with 3 individual B-scans collected at each position and 256 tomographic positions (y). Repeated volumetric scans were performed at a time interval of 6 s at baseline, during the stimulation period and during a 20-s poststimulation period to record the time course of retinal functional hyperaemia. Notably, a real-time en face OCTA display was created on the basis of a graphics processing unit (RTX 2080 Ti). During dynamic imaging, instant feedback on the quality of the acquired angiogram was provided, allowing the operators to adjust the position of the OCT module and improve the scan yield rate by ensuring that the images were stable and preventing bulk motion artefacts in the data (26).

### fOCTA data processing

In each volume, the spectral interferogram was Fourier transformed to construct the OCT structure. To extract the dynamic blood flow signals, especially in deep tissue regions, the inverse signal-to-noise ratio and decorrelation OCT angiography (ID-OCTA) algorithm were applied to the OCT angiogram (44). Retinal layer segmentation was performed via a graph search algorithm (45), and three laminar vascular/capillary plexuses, including the arteriole/venule/SCP, ICP, and DCP, were generated by projecting the OCTA signals within specific retinal slabs. The artefacts of the major vessels on the ICP and DCP slabs were subtracted on the basis of their intensity-normalized decorrelation values (46) to generate conventional (Fig. 3a) and functional (Figs. 3b and 4a-4b) OCTA images.

### Vascular skeleton-based dynamic registration

To further minimize the spatial misalignment caused by eye motion in 160 s dynamic imaging in vivo, a skeleton-based dynamic registration method was developed, which enables spatial registration of OCTA image sequences while preserving vascular functional hyperaemic response. First, the raw OCTA sequence of SVP was registered by optical flow method with deep neural network (47). The registered sequence of SVP was averaged and then served as a ground truth of SVP slab. Second, the raw OCTA sequence of SVP and the corresponding ground truth were binarized by Otsu threshold (26) and then skeletonized. Using the KAZE feature points [48], the skeleton of SVP sequence was registered based on the skeleton of SVP ground truth. Therefore, geometric transformation relationship for OCTA image at each timepoint can be generated. Based on the relationship, a rigid transformation is applied to the dynamic sequences of SVP, ICP and DCP. Finally, registered OCTA image sequences that preserve the dynamic information of vascular response were obtained.

### Quantification of functional rNVC

In one trial, each en face OCTA image was first binarised with a consistent Otsu threshold (Supplementary Fig. 11) (26). An annulus centred on the optic nerve head with inner and outer ring diameters of 0.6 and 1.8 mm, respectively, was selected as the region of interest (48). We define *P*(*x,y*) = 1 for pixels in this region and *P*(*x,y*) = 0 for those outside this region. The skeletons of the arterioles and venules were obtained by setting a decorrelation threshold to remove capillaries with lower values. Then, *A*(*x,y*) and *S*(*x,y*) were calculated as the number of pixels occupied by the blood vessels and the vascular skeleton, respectively. The VC (representing the vessel width) of the retinal arterioles and venules was then obtained as follows:

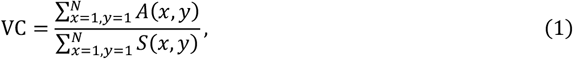

where *N* is the width/height of the square image. Each pixel on the skeleton was dilated to the corresponding VC value to generate masks for the arterioles and venules. The major vessels were subsequently subtracted from the = binarised angiograms of the ICP and DCP to remove the influence of artefacts on the quantified data. Notably, to calculate the incidence of dilation, the major vessel was first segmented on the basis of the bifurcation results, and then the mean value of ΔVC 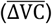 after FLS onset was used to classify single arterioles or venules as vasodilation 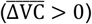 and vasoconstriction 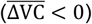 types.

For the capillary plexuses, we used VD, the percentage area occupied by the capillaries, as a substitute for VC due to the limited capillary calibre that may cause error in the calculation of VC:

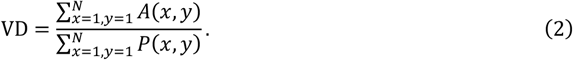

Since the decorrelation value in OCTA is positively correlated with flow velocity (49), we used decorrelation-based indices to quantify blood flow. For each vascular plexus, the decorrelation image was masked with the corresponding binarized angiogram, and the FV and RBF were calculated as the average and sum of the decorrelation values *D*(*x,y*), respectively:

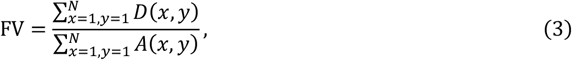

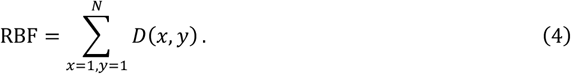

Following these formulas, the baseline and FLS-evoked VC, VD, FV, and RBF were calculated. The percentage changes in these variables, ΔVC, ΔVD, ΔFV, and ΔRBF, were used to characterize retinal functional hyperaemia (27) and were calculated as

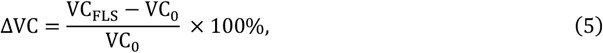

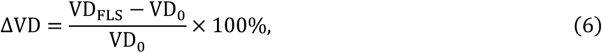

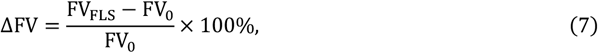

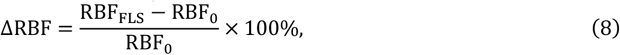

where variables with the subscript FLS refer to the peak value of the index after FLS onset, and VC_0_,VD_0_,FV_0_,and RBF_0_ denote the mean baseline values of the corresponding index before FLS. Notably, when the functional angiograms were drawn, we segmented the vasculature by the vessel bifurcation points, and the ΔRBF in each vascular segment was subsequently calculated.

The decorrelation value *D* is correlated with the blood flow speed *v* as follows:

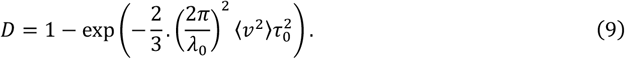

This *D* value monotonically increases with the blood velocity *v* (49), where λ_0_ is the central wavelength of the OCT probing light and τ_0_ is the correlation decay at the ms scale. Therefore, as the average of the vascular decorrelation, FV also monotonically increases with the blood flow velocity. In contrast, RBF reflects both the flow velocity and vessel size in major vessels as follows:

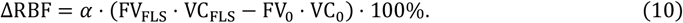

In capillaries, this relationship is written as follows:

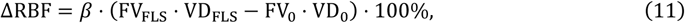

where 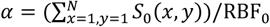 and 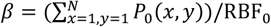 are constants for the major vessel and the capillary, respectively (derived in the Supplementary Material).

Note that ΔRBF changes with time, and we used the peak amplitude to characterize functional rNVC as follows:

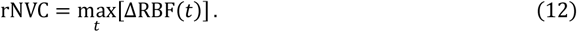

In addition, started from FLS onset, the time cost to reach the peak amplitude was calculated as the peak time of functional hyperaemia. Notably, considering the limited sampling rate, the accuracy of the peak time was improved using an intensity centroid method (50), where the Time was quantified as a weighted average centred at the peak amplitude of the functional index. Take the ΔRBF as an example:

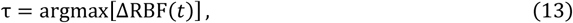

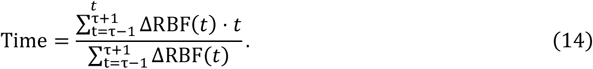

### Characterization of functional rNVC

To characterize the spatial distribution of functional hyperaemia, single capillary-resolved fOCTA images were generated as previously described (27). In brief, to improve the signal-to-noise ratio, the en face OCTA images were first averaged separately over the baseline (all 5 time points) and FLS/post-FLS (3 time points around the peak response) periods. Then, the average angiograms were binarized and skeletonized, and each vascular segment was located by removing the bifurcation points from the skeleton. The amplitude of ΔRBF in each vascular/capillary segment was used to indicate the degree of contrast in the fOCTA images.

### PD classification

We used a support vector machine to construct a hyperplane for PD classification in this study. The Euclidean distances between samples and the hyperplane are input for the receiver operating characteristic analysis and the classification accuracy was quantified with the AUROC. First, we conducted the receiver operating characteristic analysis on the basis of the outcomes of the behavioural tests (% contralateral touches in the cylinder test and the time taken to descend the pole in the pole test), the features of retinal thickness and vascular morphology (baseline VC or VD), and the functional rNVC index (the amplitude of the ΔRBF) to assess the ability of these markers to discriminate premotor PD mice from sham or aged mice. We then evaluated the classification ability of a two rNVC-variable model (ΔRBF at LDCT-Off and On). Furthermore, we leveraged rNVC indices at LDCT-Off and On in different capillary plexus to assess the classification performance of combining multiple rNVC features.

### Statistical analysis

The time courses of functional hyperaemia are expressed as mean ± s.e.m.. For comparisons between different groups, the one-tailed Mann–Whitney U test was used. Within-group differences during LDCT were assessed with the one-tailed Wilcoxon matched-pairs signed-rank test. *p* < 0.05 was considered to indicate statistical significance.

## Acknowledgments

This work is supported by the National Natural Science Foundation of China (62075189, T2293751, T2293753, 62035011); the “Pioneer” and “Leading Goose” R&D Program of Zhejiang (2023C03089).

