## Supplementary Files for "Transocular detection of premotor Parkinson’s disease via retinal capillary neurovascular coupling through functional OCT angiography"

**This PDF file includes:**

Supporting text
Figures S1 to S11
Tables S1 to S3
Legend for Movie S1

**Other supporting materials for this manuscript include the following:**

Movie S1

### Supporting Information Text

**Relationships among ΔRBF, flow velocity, and vessel size.** Retinal blood flow (RBF) is the sum of the vessel's decorrelation D, calculated as follows:

$$\mathrm{RBF} = \sum_{x=1,y=1}^{N} D(x,y).$$

Therefore, RBF changes with both vessel size (the summed area) and flow velocity (the decorrelation value).

For major vessels, since vessel calibre ($\mathrm{VC}$) is used to quantify the vessel size, we have the following equations:

$$\Delta\mathrm{RBF} = \frac{\left(\sum_{x=1,y=1}^{N} D_{FLS}(x,y) - \sum_{x=1,y=1}^{N} D_0(x,y)\right)}{\mathrm{RBF}_0} \cdot 100\%$$

$$= \frac{\left(\frac{\sum_{x=1,y=1}^{N} D_{FLS}(x,y)}{\sum_{x=1,y=1}^{N} A_{FLS}(x,y)} \cdot \sum_{x=1,y=1}^{N} A_{FLS}(x,y) - \frac{\sum_{x=1,y=1}^{N} D_0(x,y)}{\sum_{x=1,y=1}^{N} A_0(x,y)} \cdot \sum_{x=1,y=1}^{N} A_0(x,y)\right)}{\mathrm{RBF}_0} \cdot 100\%$$

$$= \frac{\left(\mathrm{FV_{FLS}} \cdot \frac{\sum_{x=1,y=1}^{N} A_{FLS}(x,y)}{\sum_{x=1,y=1}^{N} S_{FLS}(x,y)} \sum_{x=1,y=1}^{N} S_{FLS}(x,y) - \mathrm{FV_0} \cdot \frac{\sum_{x=1,y=1}^{N} A_0(x,y)}{\sum_{x=1,y=1}^{N} S_0(x,y)} \sum_{x=1,y=1}^{N} S_0(x,y)\right)}{\mathrm{RBF}_0} \cdot 100\%$$

$$= \frac{\left(\mathrm{FV_{FLS}} \cdot \mathrm{VC_{FLS}} \cdot \sum_{x=1,y=1}^{N} S_{FLS}(x,y) - \mathrm{FV_0} \cdot \mathrm{VC_0} \cdot \sum_{x=1,y=1}^{N} S_0(x,y)\right)}{\mathrm{RBF}_0} \cdot 100\%$$

Given that the vessel skeletons may exhibit little change before and after flicker light stimulation (FLS), i.e., $\sum_{x=1,y=1}^{N} S_{FLS}(x,y) \approx \sum_{x=1,y=1}^{N} S_0(x,y)$, the equation is as follows:

$$\Delta\mathrm{RBF} = \alpha \cdot (\mathrm{FV_{FLS}} \cdot \mathrm{VC_{FLS}} - \mathrm{FV_0} \cdot \mathrm{VC_0}) \cdot 100\%,$$

where $\alpha$ is a constant defined as $\alpha = (\sum_{x=1,y=1}^{N} S_0(x,y))/\mathrm{RBF}_0$.

Similarly, in the capillaries, vessel density ($\mathrm{VD}$) is used to quantify vessel size, which produces the following equation:

$$\Delta\mathrm{RBF} = \beta \cdot (\mathrm{FV_{FLS}} \cdot \mathrm{VD_{FLS}} - \mathrm{FV_0} \cdot \mathrm{VD_0}) \cdot 100\%,$$

where $\beta$ is a constant defined as $\beta = (\sum_{x=1,y=1}^{N} P_0(x,y))/\mathrm{RBF}_0$.

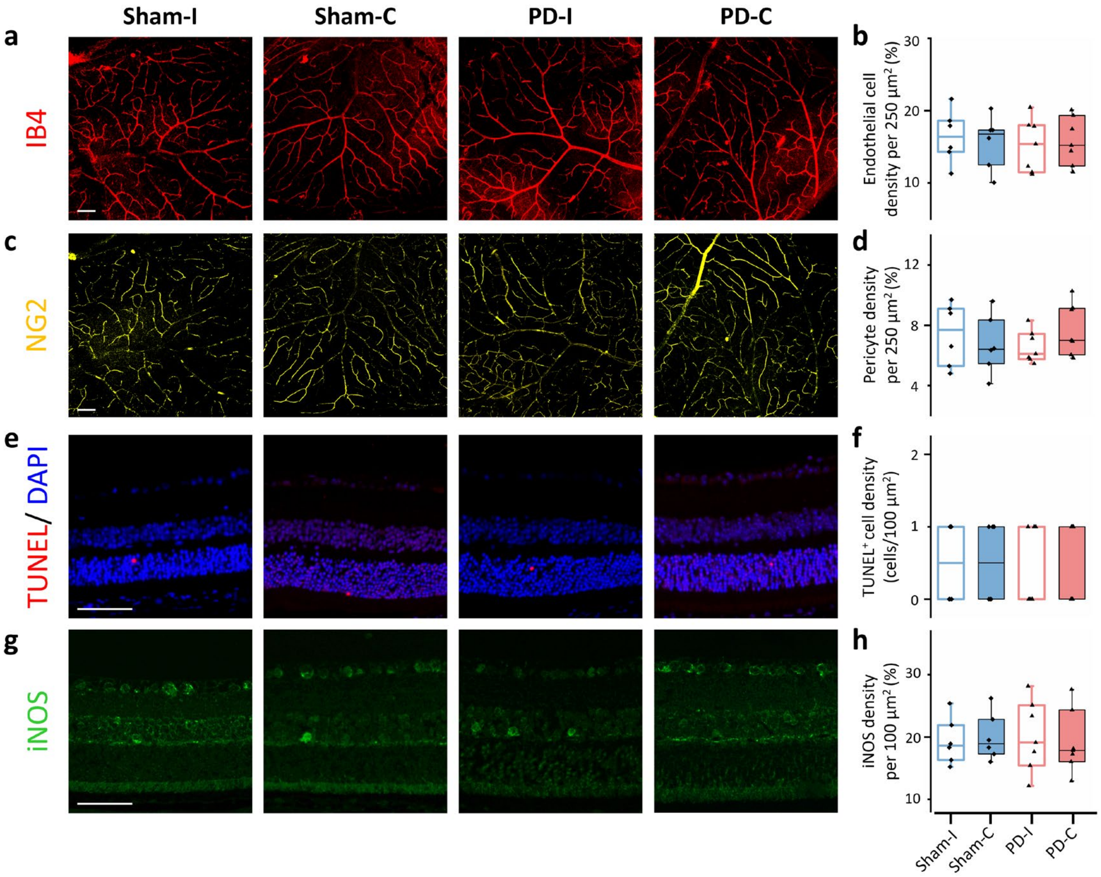


**Fig. S1.** No significant alterations in retinal vascular cells or TUNEL or iNOS expression were detected in PD mice. (a) Whole mounts of endothelial cells stained with IB4 (red) and (b) box plots of endothelial cell density. (c) Whole mounts of pericytes stained with NG2 (yellow) and (d) box plots of pericyte density. (e) Cell death as detected by TUNEL expression (red) and (f) box plots of $TUNEL^+$ cell density as a percentage. The cell nuclei are labeled with DAPI (blue). (g) Retinal sections showing iNOS expression (green) and (h) box plots of the density of iNOS-expressing cells. The pathological studies were conducted in PD group (n = 7) at 3 weeks after the 6-OHDA injections and age-matched sham group (n = 6). iNOS: inducible nitric oxide synthase; Sham-I: ipsilateral retina of the sham group; Sham-C: contralateral retina of the sham group; PD-I: ipsilateral retina of the PD group; PD-C: contralateral retina of the PD group. The scale bars represent 50 μm. * $p < 0.05$ for comparisons shown; one-tailed Mann–Whitney *U* test.

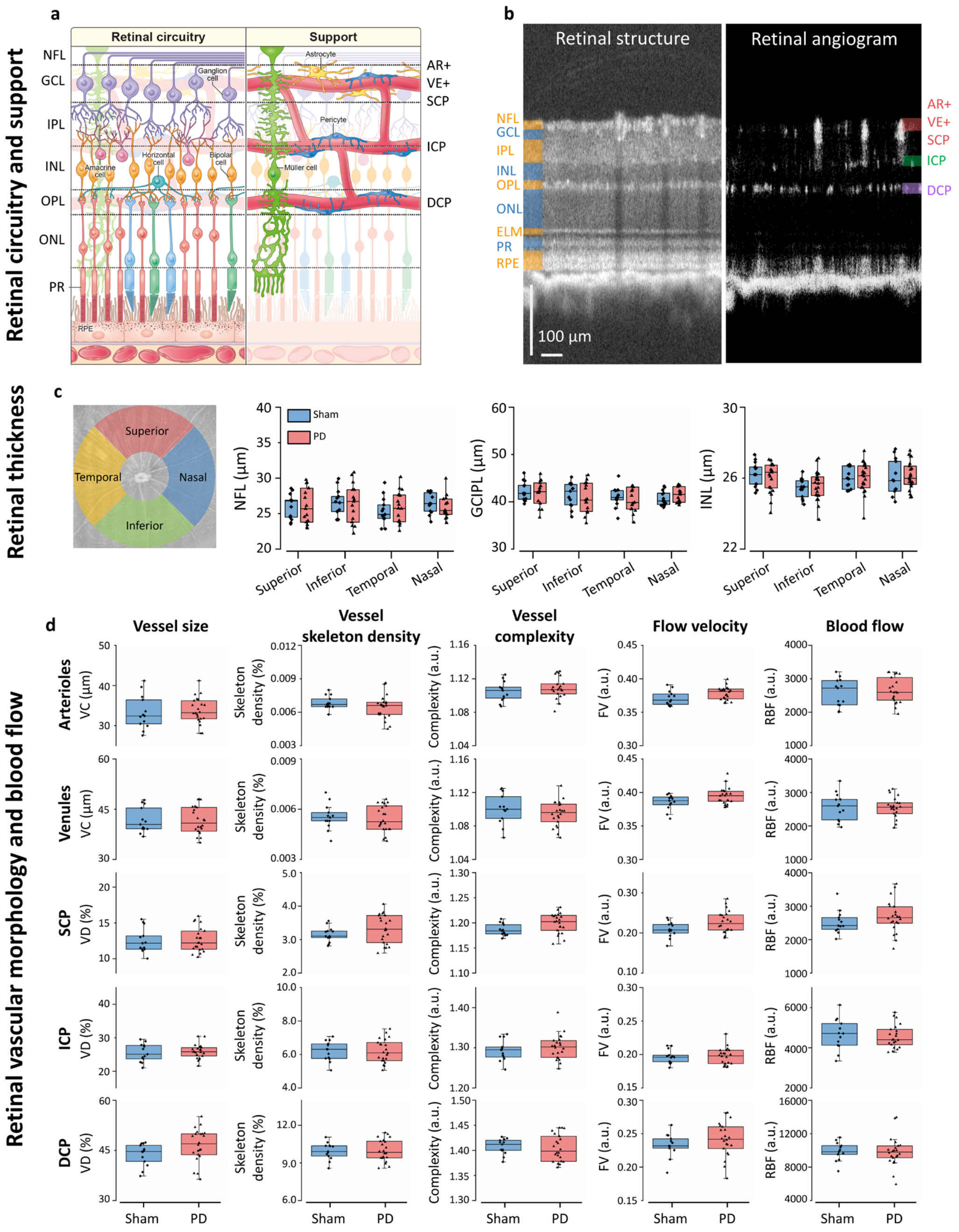

**Fig. S2.** No significant alterations in retinal thickness or baseline blood vessel size or blood flow were detected in premotor PD mice. (a) Illustration of retinal circuitry and support, adapted from Grimes et al. (25). (b) Representative cross-sectional structure and angiogram of the sham retina generated by OCT and OCTA, respectively. (c) Retinal thickness (NFL, GCIPL, and INL) was measured in the four quadrants within the annulus in the sham and PD groups. (d) Retinal vessel size (VC or VD), vessel skeleton density, vessel complexity, flow velocity (FV), and blood flow (RBF) of arterioles, venules, SCP, ICP, and DCP in the sham and PD groups. Vascular complexity was calculated based on fractal dimension using the box-counting method. The contralateral retinas of sham (n = 13, aged 12 weeks) and PD (n = 22, aged 12 weeks) mice at 1 week after lesion induction (no motor deficits) were used. NFL: nerve fibre layer; GCL: ganglion cell layer; IPL: inner plexiform layer; INL: inner nuclear layer; OPL: outer plexiform layer; ONL: outer nuclear layer; ELM: external limiting membrane; PR: photoreceptor; RPE: retinal pigment epithelium; GCIPL: ganglion cell-inner plexiform layer; AR: arteriole; VE: venule; SCP: superficial capillary plexus; ICP: intermediate capillary plexus; DCP: deep capillary plexus; VC: vessel calibre; VD: vessel density; FV: flow velocity; RBF: retinal blood flow; $p < 0.05$ was considered to indicate a statistically significant difference according to the one-tailed Mann–Whitney *U* test.

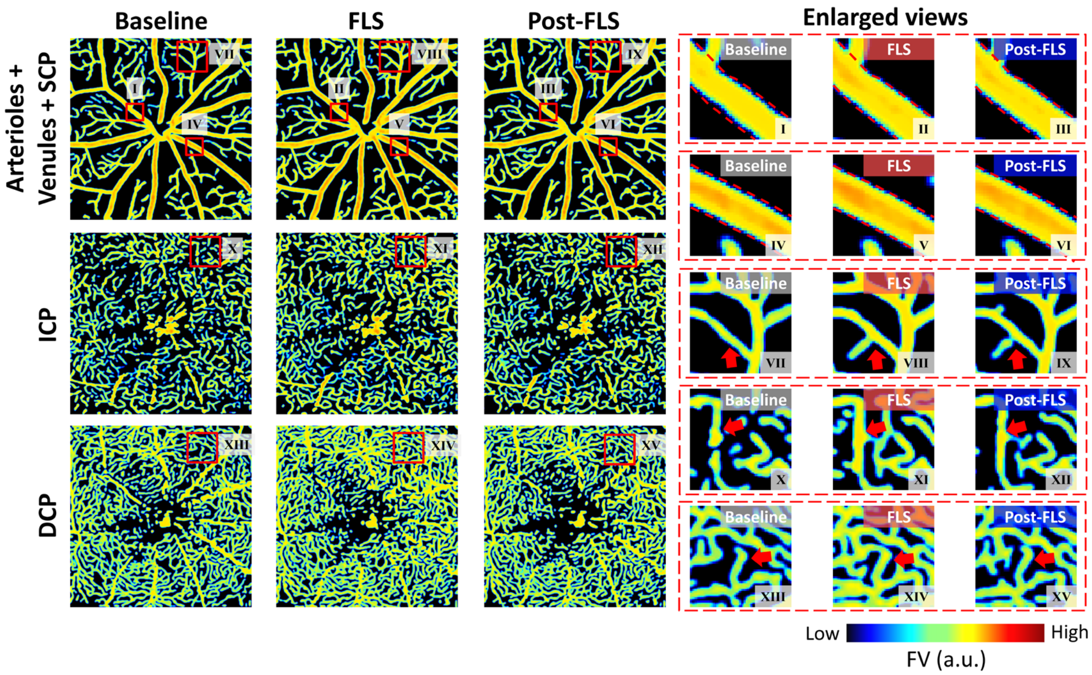


**Fig. S3.** FLS-evoked retinal functional hyperaemia in sham mice through fOCTA with single-capillary resolution. FV-encoded en face OCT angiograms (~2 mm × 2 mm) of arterioles, venules, SCP, ICP, and DCP at baseline, during FLS, and post-FLS. The contralateral retina of a sham mouse at 1 week after saline injection was used. Insets I-XV are enlarged views of the local regions indicated by red boxes. The dashed lines in I-VI mark the vessel profile during FLS, whereas the arrowheads in VII-XV indicate the obvious response in the capillary plexuses. FV: flow velocity; SCP: superficial capillary plexus; ICP: intermediate capillary plexus; DCP: deep capillary plexus; FLS: flicker light stimulation. The corresponding video is shown in Visualization 1.

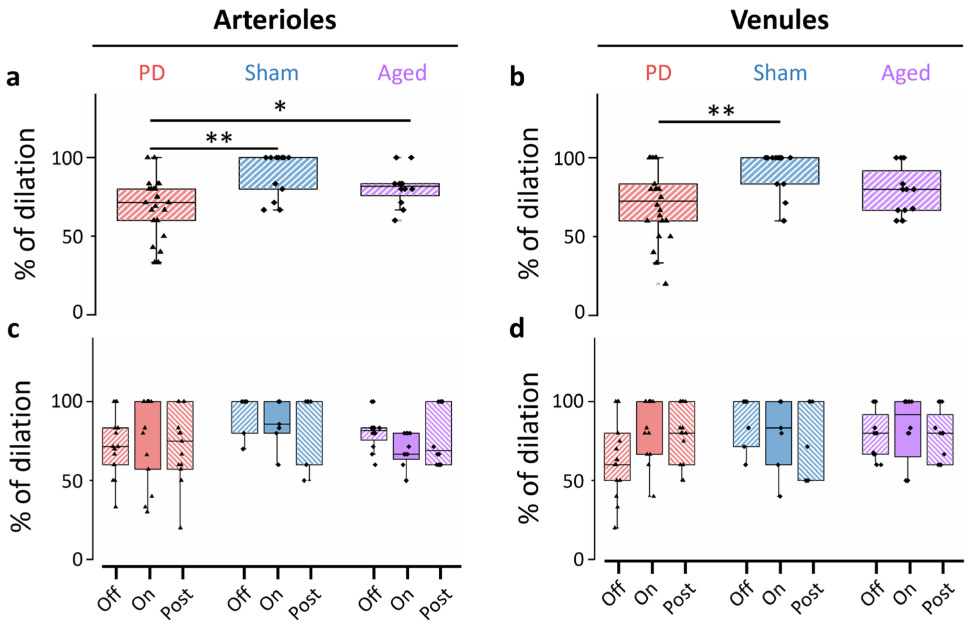


**Fig. S4.** Incidence of FLS-evoked arteriolar and venular dilation was significantly decreased in PD group at LDCT-Off and showed a trend of recovery at LDCT-On. The incidence of FLS-evoked arteriolar (a) and venular (b) dilation in the PD (n = 22), sham (n = 13) and aged (n = 12) groups at LDCT-Off. * $p < 0.05$ and ** $p < 0.01$ for comparisons shown; one-tailed Mann–Whitney *U* test. The incidences of FLS-evoked arteriolar (c) and venular (d) dilation at LDCT-Off, LDCT-On, and LDCT-Post in the PD (n = 15), sham (n = 7) and aged (n = 12) groups. $p < 0.05$ was considered to indicate a statistically significant difference according to the one-tailed Wilcoxon matched-pairs signed rank test. The incidence of dilation was defined as the percentage of dilated vessels (after FLS onset, the mean ΔVC > 0) among all vessels. The retinas contralateral to the lesion side of PD (aged 12 weeks) and sham (aged 12 weeks) mice at 1 week after lesion induction (without motor deficits) and the retinas of aged mice (aged 44 weeks) were used. LDCT: levodopa challenge test; ΔVC: percentage change in vessel calibre.

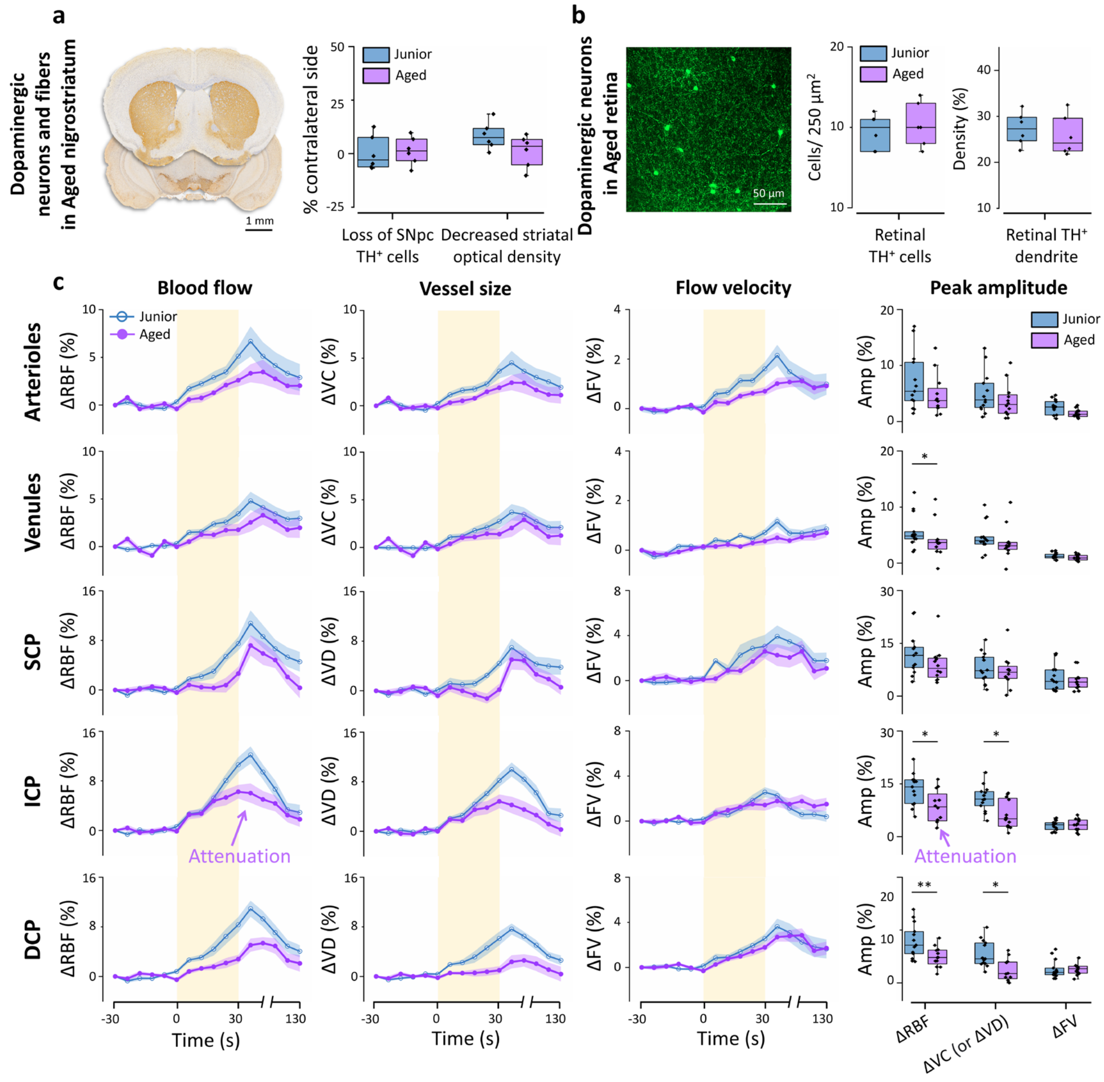


**Fig. S5.** The functional rNVC signals were attenuated in the aged group without significant loss of DAergic neurons. (a) Representative coronal brain sections stained for TH (yellow) showing the SNpc and striatum in aged mice. Box plots of SNpc TH⁺ cell loss and the decreased striatal optical density in the junior and aged groups. (b) Retinal whole mounts of TH⁺ DAergic neurons (green) from the aged group. Box plots of retinal TH⁺ cell body and dendrite density in the junior and aged groups. Junior group: n = 6, aged 12 weeks. Aged group: n = 6, aged 44 weeks. (c) Time courses of blood flow (ΔRBF), vessel size (ΔVC or ΔVD), and blood flow velocity (ΔFV) in the junior (n = 13) and aged (n = 12) groups. The yellow shaded regions indicate the period over which FLS was performed. Data are mean ± s.e.m.. The corresponding peak amplitudes of ΔRBF, ΔVC or ΔVD and ΔFV in the arterioles, venules, SCP, ICP, and DCP are plotted. The results of the junior group were obtained from the contralateral retina of sham mice (aged 12 weeks) at 1 week after saline injection. TH: tyrosine hydroxylase, an enzyme involved in the synthesis of DA that identifies DAergic cells; SNpc: substantia nigra pars compacta; SCP: superficial capillary plexus; ICP: intermediate capillary plexus; DCP: deep capillary plexus; ΔRBF, ΔVC, ΔVD, and ΔFV: percentage changes in retinal blood flow, vessel calibre, vessel density, and flow velocity, respectively. FLS: flicker light stimulation. * $p < 0.05$ and ** $p < 0.01$ for comparisons shown; one-tailed Mann–Whitney *U* test.

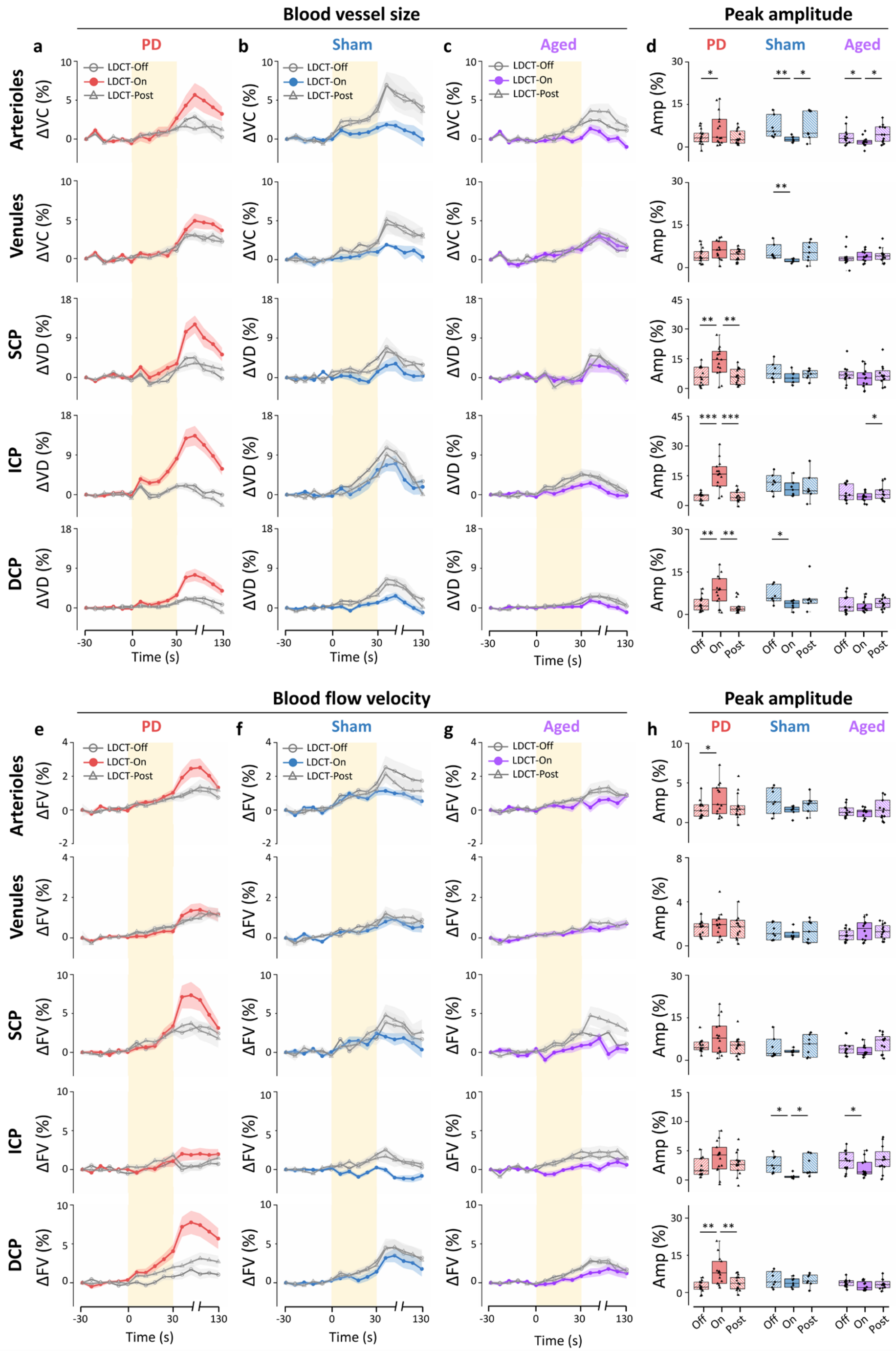


**Fig. S6.** The peak amplitudes of retinal ΔVC (or ΔVD) and ΔFV are recovered during the LDCT in PD mice, whereas they are not recovered in healthy control mice. Time courses of blood vessel size (ΔVC or ΔVD) in PD (a), sham (b), and aged (c) mice at LDCT-Off, LDCT-On, and LDCT-Post. (d) Box plots of the peak amplitudes of ΔVC or ΔVD. Time courses of blood flow velocity (ΔFV) in PD (e), sham (f), and aged (g) mice at LDCT-Off, LDCT-On, and LDCT-Post. (h) Box plots of the corresponding peak amplitudes of ΔFV. The retinas contralateral to the lesion side in PD (n = 15, aged 12 weeks) and sham (n = 7, aged 12 weeks) mice at 1 week after lesion induction (without motor deficits) and the retinas of aged mice (n = 12, aged 44 weeks) were used. The yellow shaded region of the time course indicates the period of flicker light stimulation. Data in the time courses are mean ± s.e.m.. LDCT: levodopa challenge test; ΔVC, ΔVD and ΔFV: percentage changes in vessel calibre, vessel density, and flow velocity, respectively; SCP: superficial capillary plexus; ICP: intermediate capillary plexus; DCP: deep capillary plexus. * $p < 0.05$, ** $p < 0.01$, *** $p < 0.001$, LDCT-On vs. LDCT-Off or LDCT-Post in the same group, one-tailed Wilcoxon matched-pairs signed rank test. † $p < 0.05$, PD at LDCT-On vs. Sham at LDCT-Off, one-tailed Mann–Whitney *U* test.

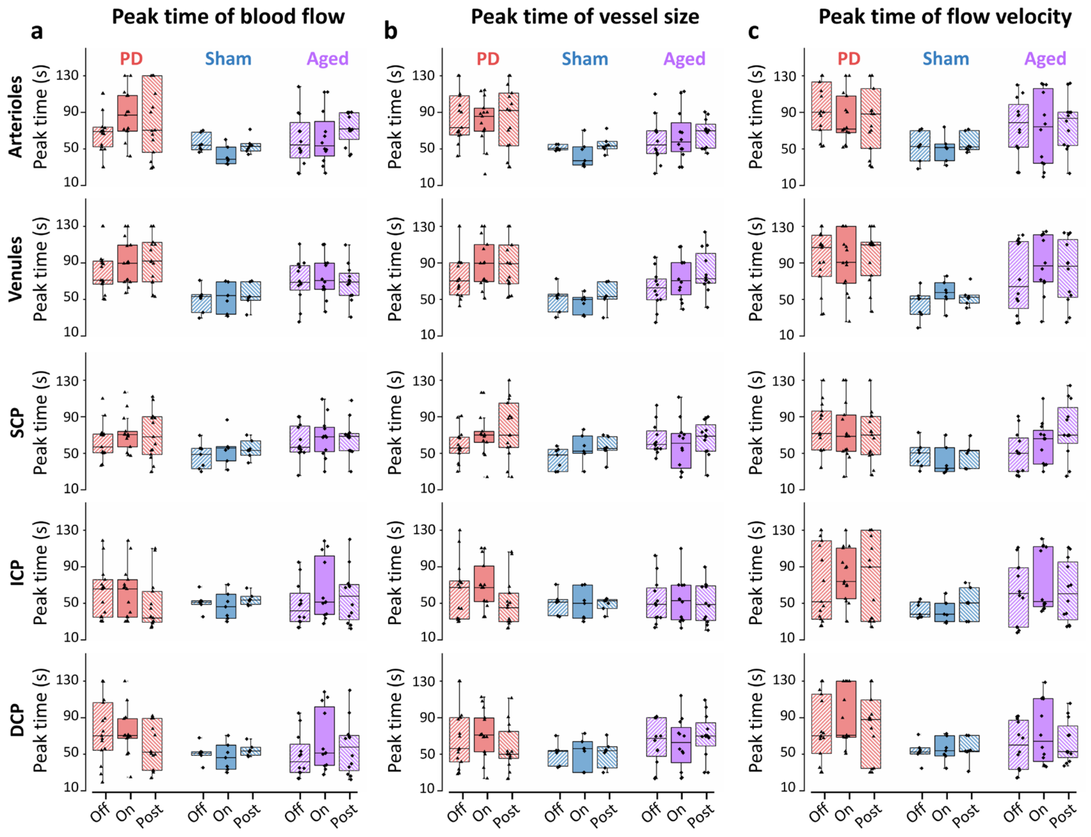


**Fig. S7.** No significant change in peak time was observed in PD, sham, or aged mice during LDCT. The box plots show the peak time of blood flow (ΔRBF) (a), vessel size (ΔVC or ΔVD) (b), and flow velocity (ΔFV) (c) of the arterioles, venules, SCP, ICP, and DCP in PD, sham, and aged mice at LDCT-Off, LDCT-On, and LDCT-Post. The retinas contralateral to the lesion side of PD (n = 15, aged 12 weeks) and sham (n = 7, aged 12 weeks) mice at 1 week after lesion induction (without motor deficits) and the retinas of aged mice (n = 12, aged 44 weeks) were used. SCP: superficial capillary plexus; ICP: intermediate capillary plexus; DCP: deep capillary plexus; ΔRBF, ΔVC, ΔVD, and ΔFV: percentage changes in retinal blood flow, vessel calibre, vessel density, and flow velocity, respectively. $p < 0.05$ was considered to indicate a statistically significant difference according to the one-tailed Wilcoxon matched-pairs signed rank test.

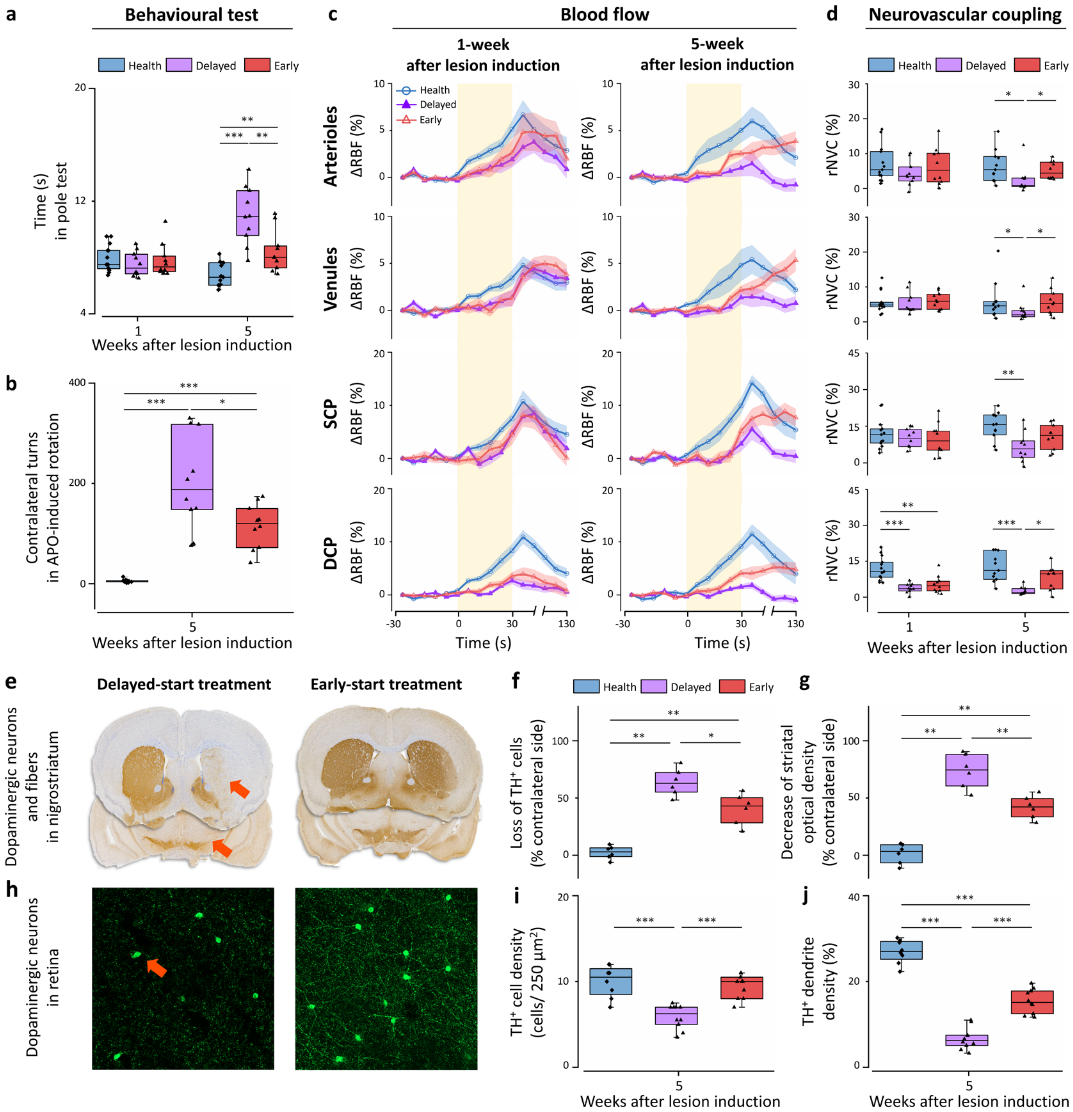


**Fig. S8.** fOCTA-rNVC allows for the early detection of premotor PD and facilitates prompt treatment with superior recovery. The pole test (a) in PD mice with early-start and delayed-start treatment and aged-matched healthy sham mice at 1-week and 5-week after lesion induction. The APO-induced rotation test (b) was performed 5-week post-lesion to confirm the damage of DAergic system in the PD model. APO: apomorphine. (c) The ΔRBF time courses of arterioles, venules, SCP, and DCP. Data are mean ± s.e.m.. The yellow shaded regions indicate the period over which FLS was applied. (d) Box plots of the corresponding rNVC indices (peak amplitude of the ΔRBF time courses). (e) Representative coronal brain sections stained for TH (yellow) showing the SNpc and striatum in PD mice with delayed-start and early-start treatments, and the corresponding statistical results of SNpc $TH^+$ cell loss (f) and the decreased striatal optical density (g) at 5 weeks after surgery. (h) Whole-mount images of $TH^+$ DAergic neuronal fibres and cell bodies (green) in the retinas of PD mice with delayed-start and early-start treatments, and the corresponding statistical results of the number of $TH^+$ cell bodies per 250 $\mu m^2$ (i) and dendrite density (j) at 5 weeks after surgery. The red triangles in (e) and (h) indicate the obvious loss of DAergic neurons or dendrites. Rasagiline, as a potential neuroprotective drug in PD, was used for the treatment. TH: tyrosine hydroxylase, an enzyme involved in the synthesis of DA and used to identify DAergic cells. The retinas contralateral to the lesion side in PD and aged-matched healthy sham mice were used in the analysis. PD with early-start treatment: n = 10; PD with delayed-start treatment: n = 10. ΔRBF: percentage change in retinal blood flow; SCP: superficial capillary plexus; DCP: deep capillary plexus; FLS: flicker light stimulation. * $p < 0.05$, ** $p < 0.01$, and *** $p < 0.001$ for comparisons shown; one-tailed Mann–Whitney *U* test.

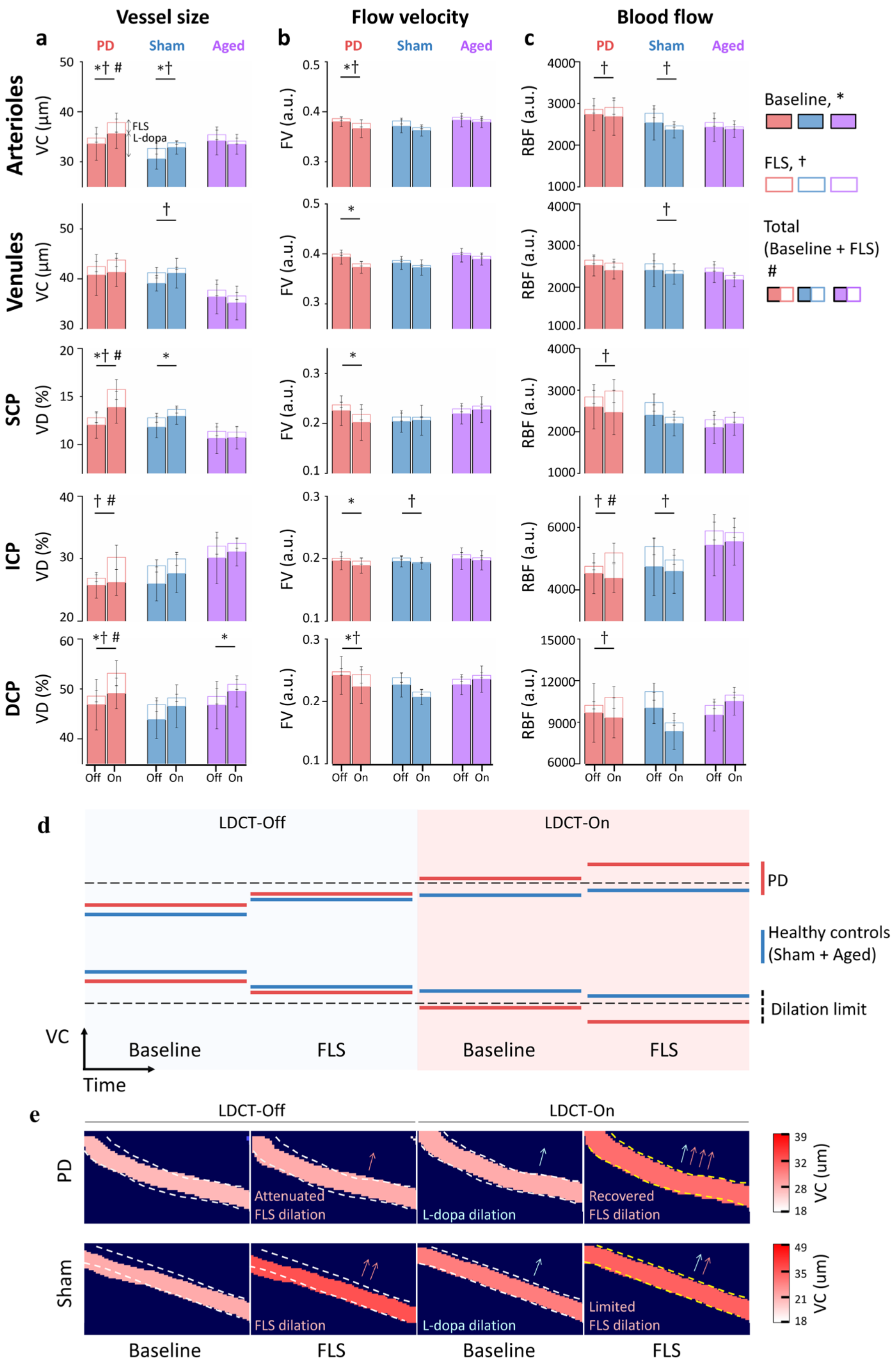


**Fig. S9.** Changes in the baseline and FLS-evoked vascular response during the LDCT. Baseline and FLS-evoked VC or VD (a), FV (b), and RBF (c) in PD, sham, and aged mice at LDCT-Off and LDCT-On. Each index value was composed of a baseline level measured before FLS and the maximal change observed during FLS. Owing to the potential vasodilation that occurs during LDCT, the VC (or VD) baseline values were elevated in the PD, sham, and aged groups at LDCT-On. Similar to the percentage changes, the absolute values of the FLS-evoked dilation indices were increased at LDCT-On than that at LDCT-Off in the PD retinas; in contrast, under FLS, these values were decreased at LDCT-On in the sham group than that at LDCT-Off. The total VC (or VD) in both the sham and aged groups did not significantly increase at LDCT-On, whereas in the PD group, these values increased significantly, surpassing the values at LDCT-Off. Besides, the FV was decreased in the PD at LDCT-On than at LDCT-Off. In addition, under FLS, the increase of FV value in the PD retinas was considerably greater at LDCT-On than at LDCT-Off, whereas the FV value increased only slightly in the sham and aged retinas at LDCT-On compared with that at LDCT-Off. Finally, the baseline RBF value was not significantly different at LDCT-On and LDCT-Off in the PD, sham, or aged groups. In contrast, under FLS, the RBF in the PD retinas increased considerably at LDCT-On compared with that at LDCT-Off, whereas the RBF increased only slightly in the sham and aged retinas at LDCT-On compared with that at LDCT-Off. The data in the bar plots are presented as the means ± SDs. (d) Schematic illustration showing the FLS-evoked VC changes at LDCT-Off and LDCT-On. Red lines: PD mice; blue lines: healthy controls (sham and aged mice); dashed lines: the possible FLS-evoked dilation limit of VC in controls. (e) Representative arterioles of PD and sham mice showing FLS-evoked VC changes at LDCT-Off and LDCT-On. The dashed lines in PD and sham mice mark the vessel profile during FLS at LDCT-On. The retinas contralateral to the lesion side of the PD (n = 15, aged 12 weeks) and sham (n = 6, aged 12 weeks) groups at 1 week after the lesion, with no motor deficits, and the retinas of the aged group (n = 12, aged 44 weeks) were used. Thus, the inhibited rNVC is most likely due to the increase in the baseline values due to levodopa-induced vasodilation, which subsequently limits the percentage changes in light-evoked hyperaemia in healthy mice. However, this overall limitation does not apply to PD mice, in which a significant increase of total value in the rNVC indices was observed at LDCT-On. VC: vessel calibre; VD: vessel density; FV: flow velocity; RBF: retinal blood flow; SCP: superficial capillary plexus; ICP: intermediate capillary plexus; DCP: deep capillary plexus; FLS: flicker light stimulation; L-dopa: levodopa; LDCT: levodopa challenge test. * $p < 0.05$ baseline at LDCT-Off vs. LDCT-On, † $p < 0.05$ FLS dilation at LDCT-Off vs. LDCT-On, # $p < 0.05$ total value (baseline + FLS dilation) at LDCT-Off vs. LDCT-On, one-tailed Wilcoxon matched-pairs signed rank test.

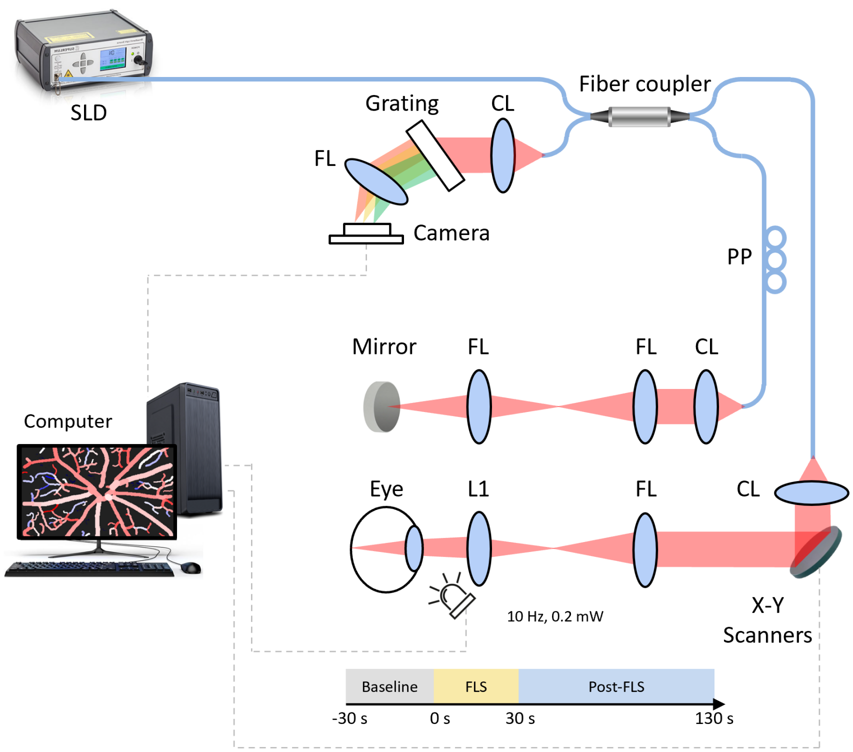


**Fig. S10.** Schematic of the fOCTA system. FLS was generated by a white light-emitting diode. The FLS power on the pupil was ~0.2 mW, with a pulse width of 50 ms and a 10 Hz repetition rate of pulse trains. Each FLS trial consisted of a 30 s baseline period followed by a 30 s FLS and a subsequent 100 s post-FLS period. FLS: flicker light stimulation; SLD: superluminescent diode; CL: collimation lens; FL: focus lens; PP: polarization paddle; L1: ocular lens.

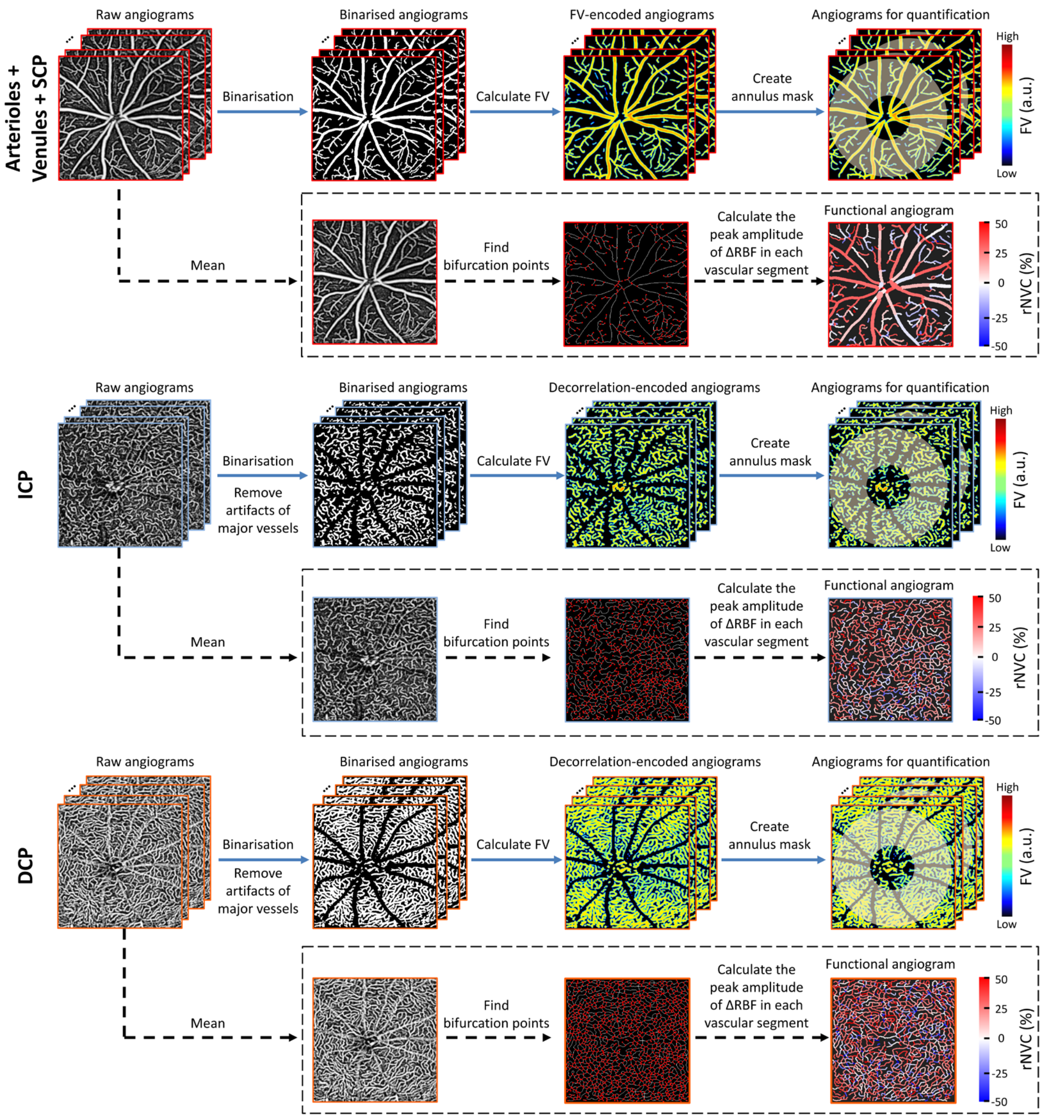


**Fig. S11.** Framework of the quantification and characterization of functional rNVC. Quantification: The raw images of each vascular plexus are binarised; then, the artefacts of major vessels are subtracted from the masks of the ICP and DCP to completely remove the influence of artefacts. FV-encoded images of arterioles, venules, and the SCP, ICP, and DCP are generated. Finally, an annulus centred on the optic nerve head with inner and outer ring diameters of 0.6 and 1.8 mm, respectively, is selected as the region of interest for hyperaemia quantification. Characterization: The raw images are first averaged to improve the signal-to-noise ratio; then, the mean angiogram is skeletonized, and each vascular segment is located by finding the bifurcation points; the mean peak amplitude of ΔRBF in each vascular/capillary segment is calculated and used as the degree of contrast of the fOCTA images. SCP: superficial capillary plexus; ICP: intermediate capillary plexus; DCP: deep capillary plexus; FV: flow velocity; ΔRBF: percentage change in retinal blood flow.

**Table S1.** Quantification of motor deficits and nigrostriatal and retinal pathology in Sham and PD mice.

| Target | Type | Time after injection | Group | Mean ± s.e.m. | *p* value |
|---|---|---|---|---|---|
| Behavioural tests | Cylinder test (%) | 1 wk | Sham (n = 13) | 50.3 ± 0.8 | |
| | | | PD (n = 22) | 49.7 ± 0.6 | > 0.05 (vs. Sham) |
| | | 2 wk | Sham (n = 6) | 50.8 ± 0.7 | |
| | | | PD (n = 7) | 49.6 ± 0.7 | > 0.05 (vs. Sham) |
| | | 3 wk | Sham (n = 6) | 50.3 ± 0.8 | |
| | | | PD (n = 7) | 46.0 ± 1.2 | **** < 0.01 (vs. Sham)** |
| | Pole test (s) | 1 wk | Sham (n = 13) | 7.9 ± 0.3 | |
| | | | PD (n = 22) | 7.6 ± 0.2 | > 0.05 (vs. Sham) |
| | | 2 wk | Sham (n = 6) | 7.0 ± 0.4 | |
| | | | PD (n = 7) | 7.2 ± 0.2 | > 0.05 (vs. Sham) |
| | | 3 wk | Sham (n = 6) | 6.6 ± 0.3 | |
| | | | PD (n = 7) | 7.8 ± 0.3 | *** < 0.05 (vs. Sham)** |
| | APO-induced rotations (turns) | 3 wk | Sham (n = 13) | 2.5 ± 0.4 | |
| | | | PD (n = 22) | 53.4 ± 5.1 | *** < 0.001 (vs. Sham)** |
| Nigrostriatal pathology | SNpc cells (%) | 1 wk | Sham (n = 6) | 0.2 ± 3.3 | |
| | | | PD (n = 7) | 14.1 ± 3.2 | *** < 0.05 (vs. Sham)** |
| | | 2 wk | Sham (n = 6) | 3.0 ± 2.9 | |
| | | | PD (n = 7) | 51.6 ± 4.0 | *** < 0.05 (vs. Sham)** |
| | | 3 wk | Sham (n = 6) | 1.7 ± 3.0 | |
| | | | PD (n = 7) | 58.2 ± 8.5 | **** < 0.01 (vs. Sham)** |
| | Striatum fibres (%) | 1 wk | Sham (n = 6) | 8.3 ± 2.6 | |
| | | | PD (n = 7) | 26.3 ± 4.5 | **** < 0.01 (vs. Sham)** |
| | | 2 wk | Sham (n = 6) | 3.1 ± 1.5 | |
| | | | PD (n = 7) | 53.8 ± 4.2 | *** < 0.05 (vs. Sham)** |
| | | 3 wk | Sham (n = 6) | 2.4 ± 2.6 | |
| | | | PD (n = 7) | 60.5 ± 5.7 | **** < 0.01 (vs. Sham)** |
| Retinal pathology | $TH^+$ cell density (cells/250 $\mu m^2$) | 1 wk | Sham-C (n = 6) | 9.5 ± 0.9 | |
| | | | Sham-I (n = 6) | 10.0 ± 0.9 | > 0.05 (vs. Sham-C) |
| | | | PD-C (n = 7) | 8.0 ± 0.9 | > 0.05 (vs. Sham-C) |
| | | | PD-I (n = 7) | 8.4 ± 0.9 | > 0.05 (vs. Sham-C) |
| | | 2 wk | Sham-C (n = 6) | 9.2 ± 0.7 | |
| | | | Sham-I (n = 6) | 9.0 ± 0.9 | > 0.05 (vs. Sham-C) |
| | | | PD-C (n = 7) | 5.7 ± 0.8 | *** < 0.05 (vs. Sham-C)** |
| | | | PD-I (n = 7) | 8.6 ± 0.9 | > 0.05 (vs. Sham-C) |
| | | 3 wk | Sham-C (n = 6) | 9.2 ± 0.5 | |
| | | | Sham-I (n = 6) | 9.2 ± 0.7 | > 0.05 (vs. Sham-C) |
| | | | PD-C (n = 7) | 4.3 ± 0.6 | **** < 0.01 (vs. Sham-C)** |
| | | | PD-I (n = 7) | 8.1 ± 0.6 | > 0.05 (vs. Sham-C) |

| | | | | | |
|---|---|---|---|---|---|
| | TH⁺ dendrite density (%) | 1 wk | Sham-C (n = 6) | 29.3 ± 1.5 | |
| | | | Sham-I (n = 6) | 26.9 ± 1.0 | > 0.05 (vs. Sham-C) |
| | | | PD-C (n = 7) | 20.9 ± 1.3 | *** < 0.05 (vs. Sham-C)** |
| | | | PD-I (n = 7) | 26.3 ± 1.0 | > 0.05 (vs. Sham-C) |
| | | 2 wk | Sham-C (n = 6) | 27.2 ± 1.3 | |
| | | | Sham-I (n = 6) | 27.6 ± 1.5 | > 0.05 (vs. Sham-C) |
| | | | PD-C (n = 7) | 15.0 ± 1.6 | **** < 0.01 (vs. Sham-C)** |
| | | | PD-I (n = 7) | 25.5 ± 1.1 | > 0.05 (vs. Sham-C) |
| | | 3 wk | Sham-C (n = 6) | 28.3 ± 1.9 | |
| | | | Sham-I (n = 6) | 29.0 ± 1.3 | > 0.05 (vs. Sham-C) |
| | | | PD-C (n = 7) | 5.1 ± 0.6 | **** < 0.01 (vs. Sham-C)** |
| | | | PD-I (n = 7) | 26.4 ± 1.5 | > 0.05 (vs. Sham-C) |

One-tailed Mann–Whitney *U* test.
Sham-I and Sham-C: ipsilateral and contralateral retinas of Sham mice, respectively; PD-I and PD-C: ipsilateral and contralateral retinas of PD mice, respectively.

**Table S2.** Peak amplitude and peak time of the FLS-evoked retinal vascular response in Sham (n = 13) and PD (n = 22) mice at 1 week after the injection.

| Vessel type | Functional index | Index property | Group | Mean ± s.e.m. | *p* value (vs. Sham) |
|---|---|---|---|---|---|
| Arterioles | ΔRBF | Peak amplitude (%) | Sham | 7.1 ± 1.1 | |
| | | | PD | 3.9 ± 0.6 | *** < 0.05** |
| | | Peak time (s) | Sham | 48.4 ± 3.4 | |
| | | | PD | 73.6 ± 5.4 | ***** < 0.001** |
| | ΔVC | Peak amplitude (%) | Sham | 5.1 ± 1.1 | |
| | | | PD | 3.0 ± 0.5 | > 0.05 |
| | | Peak time (s) | Sham | 44.2 ± 2.7 | |
| | | | PD | 86.3 ± 6.3 | ***** < 0.001** |
| | ΔFV | Peak amplitude (%) | Sham | 2.5 ± 0.4 | |
| | | | PD | 1.5 ± 0.2 | > 0.05 |
| | | Peak time (s) | Sham | 46.3 ± 3.9 | |
| | | | PD | 91.8 ± 6.7 | ***** < 0.001** |
| Venules | ΔRBF | Peak amplitude (%) | Sham | 5.7 ± 0.8 | |
| | | | PD | 4.4 ± 0.6 | > 0.05 |
| | | Peak time (s) | Sham | 47.6 ± 3.3 | |
| | | | PD | 81.2 ± 5.3 | ***** < 0.001** |
| | ΔVC | Peak amplitude (%) | Sham | 4.7 ± 0.8 | |
| | | | PD | 3.9 ± 0.5 | > 0.05 |
| | | Peak time (s) | Sham | 47.7 ± 3.5 | |
| | | | PD | 82.6 ± 6.5 | ***** < 0.001** |
| | ΔFV | Peak amplitude (%) | Sham | 1.3 ± 0.1 | |
| | | | PD | 1.4 ± 0.1 | > 0.05 |
| | | Peak time (s) | Sham | 46.3 ± 3.3 | |
| | | | PD | 90.8 ± 6.4 | ***** < 0.001** |
| SCP | ΔRBF | Peak amplitude (%) | Sham | 12.0 ± 1.7 | |
| | | | PD | 7.5 ± 1.1 | *** < 0.05** |
| | | Peak time (s) | Sham | 45.9 ± 3.2 | |
| | | | PD | 69.3 ± 4.8 | **** < 0.01** |
| | ΔVD | Peak amplitude (%) | Sham | 7.9 ± 1.2 | |
| | | | PD | 5.3 ± 0.9 | > 0.05 |
| | | Peak time (s) | Sham | 44.3 ± 3.1 | |
| | | | PD | 61.5 ± 4.2 | **** < 0.01** |
| | ΔFV | Peak amplitude (%) | Sham | 5.1 ± 1.0 | |
| | | | PD | 4.5 ± 0.5 | > 0.05 |
| | | Peak time (s) | Sham | 44.7 ± 3.4 | |
| | | | PD | 80.6 ± 6.1 | ***** < 0.001** |
| ICP | ΔRBF | Peak amplitude (%) | Sham | 13.4 ± 1.3 | |
| | | | PD | 5.2 ± 0.5 | ***** < 0.001** |
| | | Peak time (s) | Sham | 46.4 ± 2.9 | |
| | | | PD | 61.3 ± 6.4 | > 0.05 |
| | ΔVD | Peak amplitude (%) | Sham | 10.9 ± 1.0 | |
| | | | PD | 4.1 ± 0.4 | ***** < 0.001** |
| | | Peak time (s) | Sham | 46.9 ± 3.1 | |
| | | | PD | 62.8 ± 6.1 | > 0.05 |
| | ΔFV | Peak amplitude (%) | Sham | 3.1 ± 0.4 | |
| | | | PD | 2.2 ± 0.3 | > 0.05 |
| | | Peak time (s) | Sham | 37.6 ± 2.3 | |
| | | | PD | 72.8 ± 8.5 | *** < 0.05** |
| DCP | ΔRBF | Peak amplitude (%) | Sham | 11.9 ± 1.4 | |
| | | | PD | 4.8 ± 0.6 | ***** < 0.001** |
| | | Peak time (s) | Sham | 49.7 ± 2.5 | |
| | | | PD | 63.4 ± 6.9 | > 0.05 |
| | ΔVD | Peak amplitude (%) | Sham | 8.3 ± 1.0 | |
| | | | PD | 3.2 ± 0.5 | ***** < 0.001** |
| | | Peak time (s) | Sham | 49.2 ± 2.7 | |
| | | | PD | 63.3 ± 6.3 | > 0.05 |
| | ΔFV | Peak amplitude (%) | Sham | 3.9 ± 0.8 | |
| | | | PD | 2.4 ± 0.4 | > 0.05 |
| | | Peak time (s) | Sham | 47.4 ± 3.1 | |
| | | | PD | 73.4 ± 7.3 | > 0.05 |

One-tailed Mann–Whitney *U* test.

**Table S3.** Amplitude of the FLS-evoked retinal vascular response at LDCT-Off (< 0 h), LDCT-On (1 h), and LDCT-Post (36 h) in PD (n = 15), Sham (n = 7), and Aged (n = 12) mice.

| Vessel type | Functional index | Group | Imaging timepoint during LDCT | Mean ± s.e.m. (%) | *p* value |
|---|---|---|---|---|---|
| Arterioles | ΔRBF | PD | Off | 4.5 ± 0.8 | |
| | | | On | 8.9 ± 1.9 | ** < 0.01 (vs. Off) |
| | | | Post | 4.5 ± 1.0 | * < 0.05 (vs. On) |
| | | Sham | Off | 9.5 ± 2.1 | |
| | | | On | 4.1 ± 0.5 | * < 0.05 (vs. Off) |
| | | | Post | 9.5 ± 2.3 | > 0.05 (vs. On) |
| | | Aged | Off | 4.7 ± 1.0 | |
| | | | On | 2.3 ± 0.6 | > 0.05 (vs. Off) |
| | | | Post | 5.9 ± 1.3 | > 0.05 (vs. On) |
| | ΔVC | PD | Off | 3.5 ± 0.7 | |
| | | | On | 6.4 ± 1.4 | * < 0.05 (vs. Off) |
| | | | Post | 3.3 ± 0.6 | > 0.05 (vs. On) |
| | | Sham | Off | 7.0 ± 1.5 | |
| | | | On | 2.9 ± 0.4 | ** < 0.01 (vs. Off) |
| | | | Post | 7.5 ± 2.0 | * < 0.05 (vs. On) |
| | | Aged | Off | 3.7 ± 0.9 | |
| | | | On | 1.9 ± 0.5 | * < 0.05 (vs. Off) |
| | | | Post | 4.7 ± 0.9 | * < 0.05 (vs. On) |
| | ΔFV | PD | Off | 1.6 ± 0.3 | |
| | | | On | 2.9 ± 0.5 | * < 0.05 (vs. Off) |
| | | | Post | 1.9 ± 0.4 | > 0.05 (vs. On) |
| | | Sham | Off | 2.8 ± 0.6 | |
| | | | On | 1.5 ± 0.2 | > 0.05 (vs. Off) |
| | | | Post | 2.3 ± 0.4 | > 0.05 (vs. On) |
| | | Aged | Off | 1.5 ± 0.2 | |
| | | | On | 1.2 ± 0.4 | > 0.05 (vs. Off) |
| | | | Post | 1.8 ± 0.3 | > 0.05 (vs. On) |
| Venules | ΔRBF | PD | Off | 5.0 ± 0.8 | |
| | | | On | 7.5 ± 1.1 | > 0.05 (vs. Off) |
| | | | Post | 5.4 ± 0.7 | > 0.05 (vs. On) |
| | | Sham | Off | 6.5 ± 1.2 | |
| | | | On | 3.4 ± 0.2 | ** < 0.01 (vs. Off) |
| | | | Post | 6.8 ± 1.7 | > 0.05 (vs. On) |
| | | Aged | Off | 4.1 ± 0.9 | |
| | | | On | 4.6 ± 0.6 | > 0.05 (vs. Off) |
| | | | Post | 4.9 ± 0.9 | > 0.05 (vs. On) |
| | ΔVC | PD | Off | 4.2 ± 0.6 | |
| | | | On | 5.9 ± 0.8 | > 0.05 (vs. Off) |
| | | | Post | 4.4 ± 0.5 | > 0.05 (vs. On) |
| | | Sham | Off | 5.5 ± 1.0 | |
| | | | On | 2.4 ± 0.2 | ** < 0.01 (vs. Off) |
| | | | Post | 5.5 ± 1.3 | > 0.05 (vs. On) |
| | | Aged | Off | 3.6 ± 0.8 | |
| | | | On | 4.0 ± 0.5 | > 0.05 (vs. Off) |
| | | | Post | 4.3 ± 0.7 | > 0.05 (vs. On) |
| | ΔFV | PD | Off | 1.5 ± 0.2 | |
| | | | On | 1.9 ± 0.3 | > 0.05 (vs. Off) |
| | | | Post | 1.6 ± 0.3 | > 0.05 (vs. On) |
| | | Sham | Off | 1.3 ± 0.3 | |
| | | | On | 1.1 ± 0.2 | > 0.05 (vs. Off) |
| | | | Post | 1.3 ± 0.4 | > 0.05 (vs. On) |
| | | Aged | Off | 1.0 ± 0.1 | |
| | | | On | 1.4 ± 0.3 | > 0.05 (vs. Off) |
| | | | Post | 1.3 ± 0.2 | > 0.05 (vs. On) |
| SCP | ΔRBF | PD | Off | 9.2 ± 1.5 | |
| | | | On | 21.3 ± 3.1 | ** < 0.01 (vs. Off) |
| | | | Post | 8.8 ± 1.4 | ** < 0.01 (vs. On) |
| | | Sham | Off | 12.4 ± 3.1 | |
| | | | On | 7.1 ± 1.2 | > 0.05 (vs. Off) |
| | | | Post | 11.4 ± 2.1 | > 0.05 (vs. On) |
| | | Aged | Off | 8.9 ± 1.5 | |
| | | | On | 7.1 ± 1.5 | > 0.05 (vs. Off) |

| | | | | | |
|---|---|---|---|---|---|
| | | | Post | 11.5 ± 2.3 | > 0.05 (vs. On) |
| | ΔVD | PD | Off | 6.5 ± 1.2 | |
| | | | On | 13.5 ± 1.9 | **** < 0.01 (vs. Off)** |
| | | | Post | 5.9 ± 1.0 | **** < 0.01 (vs. On)** |
| | | Sham | Off | 5.5 ± 1.0 | |
| | | | On | 2.4 ± 0.2 | > 0.05 (vs. Off) |
| | | | Post | 5.5 ± 1.4 | > 0.05 (vs. On) |
| | | Aged | Off | 7.0 ± 1.3 | |
| | | | On | 5.3 ± 1.4 | > 0.05 (vs. Off) |
| | | | Post | 6.8 ± 1.5 | > 0.05 (vs. On) |
| | ΔFV | PD | Off | 5.1 ± 0.6 | |
| | | | On | 8.3 ± 1.5 | > 0.05 (vs. Off) |
| | | | Post | 5.2 ± 0.8 | > 0.05 (vs. On) |
| | | Sham | Off | 4.5 ± 1.5 | |
| | | | On | 3.1 ± 0.3 | > 0.05 (vs. Off) |
| | | | Post | 5.2 ± 1.4 | > 0.05 (vs. On) |
| | | Aged | Off | 4.5 ± 0.8 | |
| | | | On | 3.3 ± 0.5 | > 0.05 (vs. Off) |
| | | | Post | 6.1 ± 0.9 | > 0.05 (vs. On) |
| ICP | ΔRBF | PD | Off | 5.1 ± 0.6 | |
| | | | On | 18.8 ± 1.9 | ***** < 0.001 (vs. Off)** |
| | | | Post | 5.5 ± 0.7 | ***** < 0.001 (vs. On)** |
| | | Sham | Off | 13.9 ± 2.1 | |
| | | | On | 8.2 ± 1.5 | > 0.05 (vs. Off) |
| | | | Post | 11.0 ± 2.5 | > 0.05 (vs. On) |
| | | Aged | Off | 8.4 ± 1.3 | |
| | | | On | 5.3 ± 0.8 | > 0.05 (vs. Off) |
| | | | Post | 9.3 ± 1.5 | *** < 0.05 (vs. On)** |
| | ΔVD | PD | Off | 4.4 ± 0.5 | |
| | | | On | 15.4 ± 1.8 | ***** < 0.001 (vs. Off)** |
| | | | Post | 4.3 ± 0.7 | ***** < 0.001 (vs. On)** |
| | | Sham | Off | 11.4 ± 1.8 | |
| | | | On | 8.7 ± 1.6 | > 0.05 (vs. Off) |
| | | | Post | 9.4 ± 2.7 | > 0.05 (vs. On) |
| | | Aged | Off | 6.3 ± 1.2 | |
| | | | On | 4.4 ± 0.7 | > 0.05 (vs. Off) |
| | | | Post | 6.4 ± 1.0 | *** < 0.05 (vs. On)** |
| | ΔFV | PD | Off | 2.0 ± 0.4 | |
| | | | On | 3.8 ± 0.7 | > 0.05 (vs. Off) |
| | | | Post | 2.8 ± 0.5 | > 0.05 (vs. On) |
| | | Sham | Off | 2.8 ± 0.6 | |
| | | | On | 0.6 ± 0.2 | *** < 0.05 (vs. Off)** |
| | | | Post | 2.4 ± 0.7 | *** < 0.05 (vs. On)** |
| | | Aged | Off | 3.2 ± 0.5 | |
| | | | On | 2.1 ± 0.4 | *** < 0.05 (vs. Off)** |
| | | | Post | 3.6 ± 0.6 | > 0.05 (vs. On) |
| DCP | ΔRBF | PD | Off | 5.7 ± 0.7 | |
| | | | On | 15.9 ± 2.2 | ***** < 0.001 (vs. Off)** |
| | | | Post | 5.5 ± 1.1 | **** < 0.01 (vs. On)** |
| | | Sham | Off | 11.7 ± 2.3 | |
| | | | On | 7.4 ± 0.9 | > 0.05 (vs. Off) |
| | | | Post | 10.6 ± 2.7 | > 0.05 (vs. On) |
| | | Aged | Off | 7.4 ± 0.8 | |
| | | | On | 4.3 ± 0.6 | *** < 0.05 (vs. Off)** |
| | | | Post | 6.7 ± 1.1 | *** < 0.05 (vs. On)** |
| | ΔVD | PD | Off | 3.6 ± 0.6 | |
| | | | On | 8.2 ± 1.3 | **** < 0.01 (vs. Off)** |
| | | | Post | 2.7 ± 0.6 | **** < 0.01 (vs. On)** |
| | | Sham | Off | 6.8 ± 1.2 | |
| | | | On | 3.6 ± 0.6 | *** < 0.05 (vs. Off)** |
| | | | Post | 6.1 ± 2.0 | > 0.05 (vs. On) |
| | | Aged | Off | 3.9 ± 0.9 | |
| | | | On | 2.9 ± 0.6 | > 0.05 (vs. Off) |
| | | | Post | 4.0 ± 0.6 | > 0.05 (vs. On) |
| | ΔFV | PD | Off | 2.4 ± 0.5 | |
| | | | On | 8.5 ± 1.4 | **** < 0.01 (vs. Off)** |

| | | | | | |
|---|---|---|---|---|---|
| | | | Post | 3.9 ± 0.7 | **** < 0.01 (vs. On)** |
| | | Sham | Off | 5.1 ± 1.2 | |
| | | | On | 3.9 ± 0.7 | > 0.05 (vs. Off) |
| | | | Post | 4.9 ± 1.0 | > 0.05 (vs. On) |
| | | Aged | Off | 3.9 ± 0.5 | |
| | | | On | 2.6 ± 0.6 | > 0.05 (vs. Off) |
| | | | Post | 3.4 ± 0.6 | > 0.05 (vs. On) |

The contralateral retinas of Sham and PD mice at 1 week after the injection and the retinas of Aged mice were examined. One-tailed Wilcoxon matched-pairs signed rank test.

**Movie S1 (separate file)**. FLS-evoked retinal functional hyperaemia in sham mice through fOCTA with single-capillary resolution.
